# No Evidence of an Effect of a Single Mirror Visual Feedback Session on Subsequent Motor Imagery

**DOI:** 10.64898/2026.11.05.763949

**Authors:** Linn Julie Schwarz, Annette Sterr, Cornelia Kranczioch

**Author notes:** Correspondence: Linn Julie Schwarz, Cornelia Kranczioch.

## Abstract

Motor imagery (MI) and mirror therapy, based on mirror visual feedback (MVF), are recommended as adjuncts to conventional therapies for upper limb rehabilitation after stroke. Both approaches activate sensorimotor cortical networks despite the absence of overt movement, and their concurrent application has been associated with enhanced neural activation. The present study investigated whether a single session of MVF modulates subsequent kinesthetic Motor Imagery (kMI)-related neural activity. EEG data from 38 healthy participants (MVF: *n* = 18; sham: *n* = 20) were analyzed in a mixed pre-post design. Participants performed kMI before and after either an MVF or sham intervention. Analyses focused on mu-band (8-12 Hz) event-related desynchronization (ERD) and its hemispheric lateralization. Although kMI reliably elicited stronger contralateral than ipsilateral mu ERD, MVF did not significantly affect subsequent kMI-related mu ERD or its hemispheric lateralization. These findings suggest that previously reported facilitatory effects of MVF on MI may be more closely related to concurrent application than to immediate aftereffects, while potential effects of repeated MVF exposure on subsequent MI remain to be investigated. These findings provide a foundation for future studies investigating how MVF and MI can be combined to optimize upper-limb stroke rehabilitation.

## 1. Introduction

Motor Imagery (MI) and Mirror Visual Feedback (MVF) are rehabilitation approaches that can engage sensorimotor brain networks despite the absence of movement of the affected limb (Deconinck et al., 2015; Munzert et al., 2009). As such, they are particularly relevant for individuals with stroke who are unable to sufficiently activate these networks through voluntary movement due to hemiparesis (Page & Peters, 2014; Thieme et al., 2018). Consistent with this rationale, both approaches are recommended as adjuncts to conventional therapy in current stroke rehabilitation guidelines (ISWP, 2023; Kwakkel et al., 2023; NICE, 2023). Given the substantial overlap in the neural networks recruited by MI and MVF (Bello et al., 2020), understanding whether their effects can interact or even reinforce each other may provide important insights for optimizing rehabilitation strategies. However, despite growing interest in the relationship between these approaches, the extent to which MVF influences subsequent MI-related sensorimotor activation remains largely unclear.

MI refers to the mental rehearsal of a movement without its overt execution and is commonly distinguished into a visual and kinesthetic modality (vMI and kMI; Dickstein & Deutsch, 2007; Jeannerod, 1995; Munzert et al., 2015). In vMI, individuals form a mental representation of how a movement looks, whereas in kMI, the focus lies on the internal sensory experience associated with movement execution (Dickstein & Deutsch, 2007). According to the neural simulation of action theory, imagined and executed movements show a functional similarity (Jeannerod, 2001). Consistent with this view, MI is associated with event-related desynchronization (ERD) in the mu (8–12 Hz) and beta (13–30 Hz) frequency bands, reflecting the engagement of task-relevant sensorimotor areas (Neuper & Pfurtscheller, 1999; Pfurtscheller et al., 2006). Mu and beta ERD are typically strongest over sensorimotor regions contralateral to the imagined movement (McFarland et al., 2000; Yuan et al., 2010). Notably, kMI elicits more robust sensorimotor rhythm modulation than vMI, suggesting a closer correspondence to motor execution and sensorimotor network engagement (Lim et al., 2006; Neuper et al., 2005). Given its ability to engage motor-related networks without requiring overt movement, MI has been widely applied in stroke rehabilitation. Several studies report improvements in upper-limb motor function when MI is used as an adjunct to conventional therapy, although findings remain somewhat inconsistent across studies (Ietswaart et al., 2011; Langhorne et al., 2009; Liu et al., 2004; Page et al., 2005; Page et al., 2007; Zimmermann-Schlatter et al., 2008).

Similar to MI, MVF has been widely applied in stroke rehabilitation and has been associated with improvements in upper-limb motor function following stroke (Morkisch et al., 2019; Thieme et al., 2018; Yang et al., 2018; Zeng et al., 2018). In MVF, movements of the non-affected limb are reflected in a mirror positioned along the individual’s midsagittal plane while the affected limb remains hidden from view. This creates the visual illusion that the affected limb is moving synchronously with the intact limb (Deconinck et al., 2015). Despite relying primarily on visual input, MVF has been shown to engage sensorimotor brain networks associated with the non-moving limb (Bai et al., 2020; Diers et al., 2010; Diers et al., 2015; Fritzsch et al., 2014; Milde et al., 2015; Numata et al., 2013; Saleh et al., 2014; Saleh et al., 2017). Consistent with this notion, oscillatory EEG studies indicate that MVF modulates sensorimotor rhythms in the mu and beta frequency ranges. In particular, MVF is commonly associated with enhanced or more widespread mu ERD, suggesting increased recruitment of sensorimotor representations corresponding to the non-moving limb (Bartur et al., 2015; Chang et al., 2023; Lee et al., 2015; Zhuang et al., 2025). In contrast, findings in the beta range are less consistent, with reported effects reflecting changes in hemispheric distribution rather than a consistent increase in sensorimotor activation (Bartur et al., 2015; Bartur et al., 2018). Accordingly, mu ERD may provide a more consistent electrophysiological marker for examining sensorimotor network engagement across MI and MVF.

Based on substantial overlap in the neural networks engaged by MI and MVF, several authors have proposed a close conceptual relationship between both approaches (Bello et al., 2020; Deconinck et al., 2015; Thieme et al., 2018). Reflecting this view, MI has been incorporated into mirror therapy protocols (Morkisch & Dohle, 2015; Rothgangel & Braun, 2013).

Initial empirical findings support the assumption that combining MI and MVF may enhance sensorimotor network engagement beyond the effects of either intervention alone. Using transcranial magnetic stimulation, studies have shown greater increases in corticospinal excitability when MI was performed concurrently with MVF compared to MI alone in both healthy individuals and stroke survivors (Fukumura et al., 2007; Kang et al., 2011). Similarly, studies employing visually induced movement representations, such as action observation or visual feedback, reported more robust and pronounced sensorimotor rhythm desynchronization during MI compared to imagery performed without additional visual input (M. Lin et al., 2023; Ono et al., 2013). Together, these findings suggest that concurrent visual movement-related input may strengthen neural processes associated with MI.

Little is known, however, about whether the facilitating effects of combining MI and MVF depend on the simultaneous presence of visual input and MI, or whether MVF can induce changes in sensorimotor network engagement that persist beyond the intervention itself (i.e., offline MVF effects). If such offline effects exist, MVF could facilitate subsequent MI even after the visual feedback has been removed. This question is of particular practical relevance because simultaneous application of MI and MVF has been described as challenging for the patient and is therefore only recommended for advanced mirror therapy patients (Morkisch & Dohle, 2015). To the best of our knowledge, only one study has directly examined whether prior exposure to MVF influences subsequent MI neural correlates. Gómez and Braidot (2024) reported stronger EEG-based indices of MI in participants who performed MVF before MI, suggesting a potential carry-over effect of MVF. However, because in their study the experimental groups differed not only in the timing of MVF exposure relative to MI but also in their motor practice prior to MI, the specific contribution of MVF to the observed effects remains difficult to isolate. Whether MVF can systematically modulate neural correlates of subsequent MI is therefore yet to be determined.

To address this gap, the present study investigated whether a single session of MVF modulates sensorimotor EEG activity of subsequent kMI. A pre-post experimental design with two groups was employed. Participants completed two kMI sessions while EEG was recorded, one preceding (MI1) and one following (MI2) an intervention phase. In the experimental group, the intervention consisted of an MVF task in which participants observed the mirrored movements of their left upper extremity, creating the visual illusion that the right upper extremity was moving. In the sham group, the same motor task was performed using a wooden board instead of a mirror, preventing MVF while keeping motor execution identical. If MVF induces sustained changes in sensorimotor network excitability, participants exposed to MVF should exhibit stronger MI-related neural activation during the post-intervention assessment. Specifically, we hypothesized that the increase in mu ERD over sensorimotor areas during kMI from MI1 to MI2 would be greater in the MVF group than in the sham group (Hypothesis 1). Furthermore, we hypothesized that the increase in lateralization of mu ERD toward contralateral sensorimotor regions from MI1 to MI2 would be greater in the MVF group than in the sham group (Hypothesis 2).

## 2. Materials and Methods

### 2.1. Participants

A total of 42 individuals (self-reported gender: 28 female, 13 male, and 1 not stated; aged 22-32 years, *M* ± *SD*: 26.21 ± 2.44 years; highest school qualification: high school diploma for all participants) participated in the experiment. All participants reported normal or corrected-to-normal vision and were free of neurological conditions. As tested with the revised Edinburgh Handedness Inventory (Veale, 2014), all participants were right-handed. They were allowed to choose their preferred language for the experiment, with 20 participants choosing German and 22 English. Recruitment took place within the student population of the University of Oldenburg via advertisements on the university’s website and word of mouth. Participants were reimbursed with €12 per hour. Two participants were excluded: one due to incompletely recorded data, and one due to discontinuation of participation because of physical discomfort. Following EEG data preprocessing and time-frequency analysis, two additional participants were excluded because they did not exhibit measurable mu ERD over contralateral left sensorimotor electrodes during MI1. Given that mu ERD is a well-established neurophysiological correlate of MI, its absence was conservatively interpreted as insufficient task compliance. This resulted in final group sizes of 18 participants in the MVF group (13 female, 4 male, and 1 not stated, aged 22-30 years, *M* ± *SD*: 25.89 years ± 2.27) and 20 participants in the sham group (13 female, aged 22-32 years, *M* ± *SD*: 26.60 years ± 2.68). Thirteen participants reported previous MI experience (7 MVF group, 6 sham group). Prior to data collection, participants were randomly assigned to one of the two groups using a custom MATLAB script (The MathWorks Inc., 2022, version R2022a) implementing equal allocation to both groups. Participants were blinded to group allocation and were not informed about the existence of two intervention conditions (MVF vs. sham). Before the experiment, participants gave written informed consent. The experiment was approved by the Commission for Research Impact Assessment and Ethics of the Carl von Ossietzky University of Oldenburg (Drs.EK/2025/001).

### 2.2. Experimental Procedure

After providing informed consent and filling in baseline questionnaires, participants received a brief introduction to MI. Then participants completed the KVIQ-10 (Malouin et al., 2007) to assess kinesthetic and visual imagery ability and gain experience with both imagery modalities. Following EEG and EMG preparation, participants were familiarized with the finger opposition movements (see Figure 1D) and pronation/supination levels used in the experimental task according to the Berlin Mirror Therapy Protocol (Morkisch & Dohle, 2015). Throughout the experiment, participants performed all tasks as if their right upper extremity were impaired and constituted the target limb for therapy. The procedure consisted of a pre-intervention MI session (MI1), an intervention phase (MVF or sham), and a post-intervention MI session (MI2; see Figure 1A).

**Figure 1:**
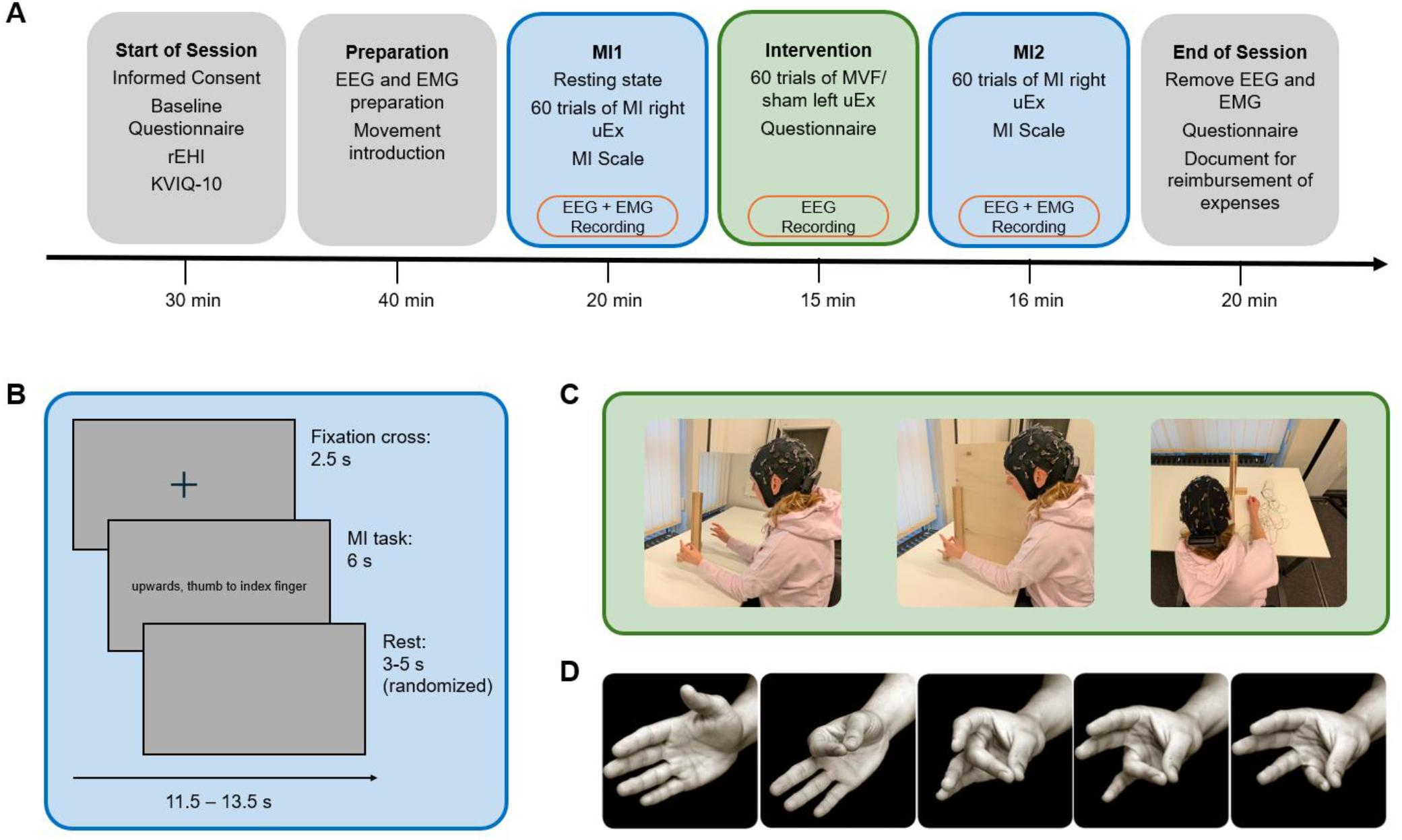
Experimental procedure and task design. (A) Overview of the experimental procedure, including the approximate duration of each phase. EEG was recorded during MI1, the intervention, and MI2, whereas EMG was recorded during MI1 and MI2 only. (B) Trial structure of the MI task. Each trial consisted of a 2.5-s fixation period, a 6-s kinesthetic MI period, and a randomized intertrial interval of 3–5 s, resulting in a total trial duration of 11.5–13.5 s. (C) Experimental setup of the intervention phase. Participants performed left upper extremity movements while viewing either a mirror (MVF; left) or a wooden panel (sham; middle) positioned in the midsagittal plane. The person depicted in panel C is one of the authors. (D) The five finger opposition movements used throughout the MI and intervention tasks according to the Berlin Mirror Therapy Protocol (Morkisch & Dohle, 2015): thumb alone, thumb to index finger, thumb to middle finger, thumb to ring finger, and thumb to little finger. Adapted from Morkisch and Dohle (2015). Reproduced with permission from Hippocampus Verlag. © 2015 Hippocampus Verlag. MI = motor imagery; MVF = mirror visual feedback; rEHI = revised Edinburgh Handedness Inventory; uEx = upper extremity.

MI1 and MI2 were conducted in an sound-attenuating, dimly lit cabin and stimuli were presented using PsychoPy (Peirce et al., 2019, version 2024.2.4). MI1 began with a 4-min resting-state recording (2 min eyes open, 2 min eyes closed). MI1 and MI2 each comprised three blocks of 20 kMI trials. A trial consisted of a 2.5-s fixation period, a 6-s imagery period, during which the movement to be imagined was displayed as a written instruction on a screen in front of the participant, and a jittered intertrial interval (range: 3-5 s with 0.5-s increments; see Figure 1B). The instructed movement was one of 15 movement combinations imagined with the right upper extremity, resulting from the combination of five finger opposition movements and three pronation/supination levels included in the Berlin Mirror Therapy Protocol (Morkisch & Dohle, 2015). Each movement appeared four times within a session. Trial order was randomized with the restriction that each movement appeared at least once but no more than twice within a block. There was a 1-min break between blocks. After every 10 trials, participants were asked to rate the perceived difficulty or ease of the preceding 10 trials using a 7-point Likert scale (1 = very hard to 7 = very easy).

Depending on group assignment, participants completed either an MVF or sham intervention. A mirror (MVF group) or wooden panel (sham group), both measuring 40 × 60 cm, was positioned upright in the midsagittal plane such that the resting right upper extremity remained out of sight (see Figure 1C). In the MVF condition, participants viewed the mirror reflection of their moving left upper extremity, creating the illusion that the right upper extremity was moving, whereas no such illusion was possible in the sham condition. Participants executed the same finger opposition and pronation/supination combinations with the left upper extremity as used in the MI task and were instructed to keep their gaze directed at the mirror or wooden panel, respectively. Participants in the MVF group were additionally instructed to focus on the reflected hand. The intervention comprised three blocks of 20 trials, with movement combinations verbally indicated by the experimenter and randomized using the same restrictions as in MI1 and MI2. A one-minute break was provided between blocks, during which participants rated their attention and tiredness for the preceding block. After the third block, participants completed a questionnaire regarding the intervention.

MI2 was identical to MI1, except that no resting-state recording was obtained. At the end of the experiment, participants filled in an additional questionnaire asking about their experience and whether they used a particular strategy for MI, and if so, which. The entire experimental session lasted approximately 2.5–3 h.

### 2.3. Data Acquisition

Data acquisition was performed with the mBrainTrain recording software (mBrainTrain, Belgrade, Serbia, version 5.0.1.0), which simultaneously recorded EEG and EMG data. EEG signals were acquired using a wireless Smarting Pro amplifier (mBrainTrain, Belgrade, Serbia) mounted onto a custom EEG cap (EasyCap GmbH, Herrsching, Germany). The cap was equipped with 32 passive wet electrodes (Ag/AgCl sintered) at the nominal 10–20 sites Fp1, Fp2, F3, F4, C3, C4, P3, P4, O1, O2, F7, F8, T7, T8, P7, P8, Fz, Cz, Pz, POz, FC1, FC2, CP1, CP2, FC5, FC6, CP5, CP6, FT9, FT10, TP9, and TP10. Due to a systematic cap displacement, the actual electrode locations differed from these nominal positions (cf. section 2.6). The reference electrode was placed at electrode site FCz. Sampling rate was set to 250 Hz, and electrode impedance was kept below 20 kΩ.

Surface EMG was simultaneously recorded from the right musculus flexor digitorum superficialis, musculus pronator teres, and musculus adductor pollicis using Ag/AgCl electrodes placed over the muscle belly and proximal base. Clavicle electrodes served as reference and ground. EMG data were recorded using a BrainAmp amplifier system (BrainProducts, Gilching, Germany). EMG electrode impedances were maintained below 100 kΩ before data acquisition and sampling rate was set to 1000 Hz.

EEG, EMG, and event markers were synchronized using the Lab Streaming Layer (LSL; Kothe et al., 2025). EEG was recorded during MI1, the intervention, and MI2, whereas EMG was recorded during MI1 and MI2 only.

### 2.4. Behavioral Analysis

Behavioral data comprised KVIQ-10 ratings, MI difficulty ratings collected during MI1 and MI2, attention and tiredness ratings collected during the intervention, and responses to post-intervention and post-experimental questionnaires. The attention item referred to attentive observation of the arm in the mirror in the MVF group and of the wooden panel in the sham group. The post-intervention and post-experimental questionnaires assessed subjective intervention experience and MI strategies. Separate kinesthetic and visual KVIQ-10 scores were calculated for each participant. To assess within-session changes in MI difficulty, ratings were averaged separately for each participant across the first and last three rating prompts of each session. Attention and tiredness ratings were retained separately for each intervention block. Questionnaire items administered in both groups were used for group comparisons, whereas items administered exclusively to the MVF group were analyzed descriptively.

### 2.5. EMG Analysis

EMG data were processed in MATLAB (The MathWorks Inc., 2022, version R2022b) using EEGLAB (Delorme & Makeig, 2004, version 2025.0.0). A 1-Hz high-pass Hamming-windowed finite impulse response (FIR) filter was applied to the EMG data (transition bandwidth = 2 Hz, order = 1650). Noise removal was performed using wavelet denoising (wavelet signal denoiser toolbox, The MathWorks Inc., 2022, version R2022b) with a Daubechies 4 (dB4) wavelet. Data were segmented from −1000 to 4500 ms relative to the onset of the MI cue, i.e., the presentation of the finger opposition movement and pronation/supination instruction. For each trial the standard deviation and the 250-samples centered moving standard deviation were calculated. Trials were classified as containing movement artifacts and excluded if the moving standard deviation exceeded the trial-specific standard deviation by a factor of 2.5 at any time point (*M* ± *SD*: 0.62 ± 1.20 trials, range: 0-7 of 60 trials per dataset^1^).

### 2.6. EEG Analysis

EEG data were processed in MATLAB (The MathWorks Inc., 2022, version R2022b) using EEGLAB (Delorme & Makeig, 2004, version 2025.0.0). An unintended but systematic posterior displacement of the EEG cap required an adjustment of channel coordinates prior to analysis. To this end, electrode positions were digitized (Xensor electrode digitizer, ANT Neuro, The Netherlands, version 5.0.0) for both the intended and actual cap placement. Based on this information channel coordinates were updated in the channels configuration file (for details, see Supplementary Figure S1). Analysis channels were selected based on their spatial proximity to the intended sensorimotor target regions (cf. Figure 2B). Throughout the manuscript, these channels are referred to according to their approximate functional location (central and centro-parietal). ICA was applied to the concatenated data of MI1, Intervention and MI2. Only MI1 and MI2 were included in subsequent analyses. ChatGPT (OpenAI, accessed October 2025 to July 2026) assisted with coding selected analysis steps in MATLAB. AI-generated code was critically reviewed, tested, and revised where necessary.

**Figure 2:**
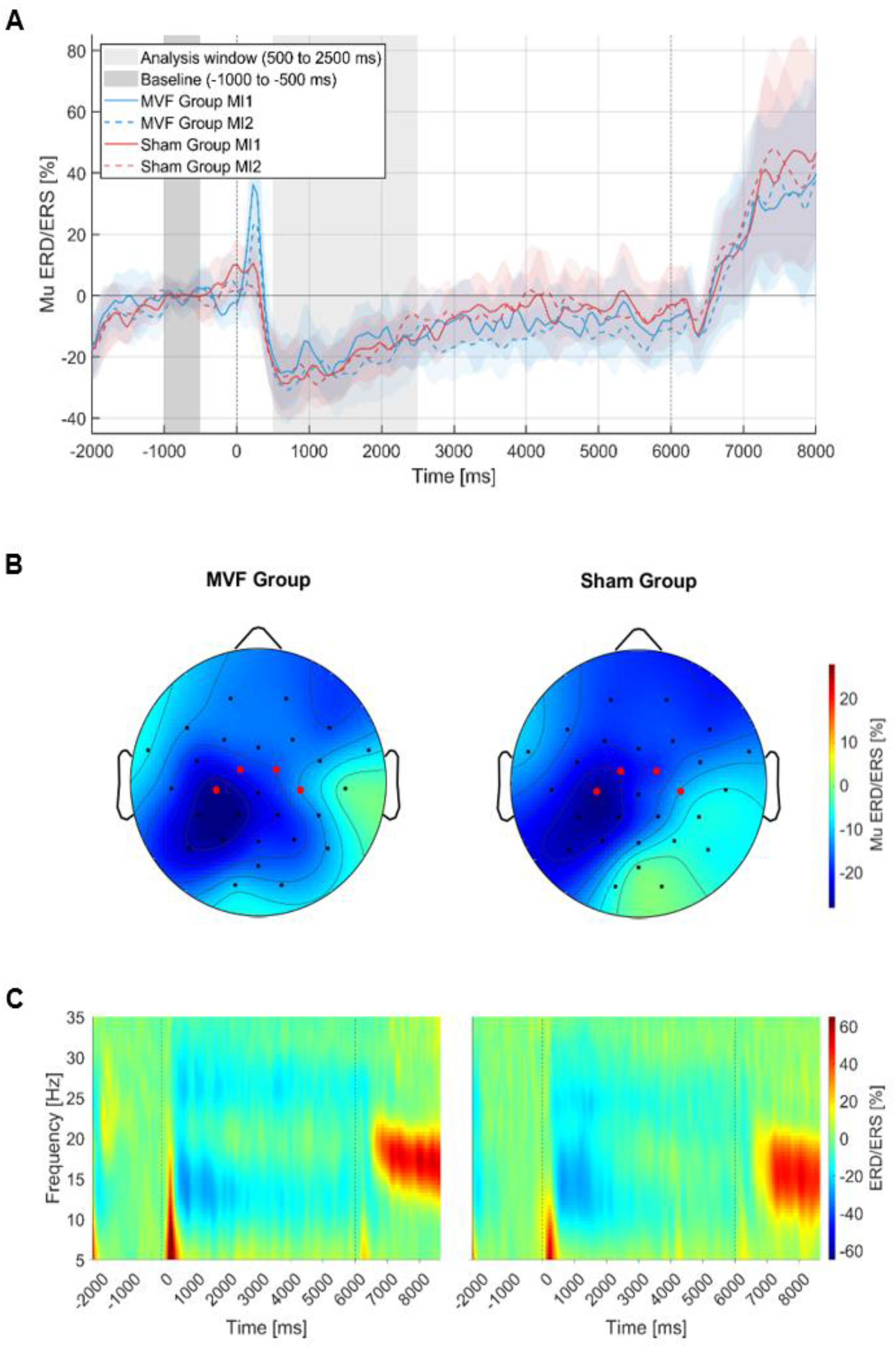
Descriptive mu event-related desynchronization/synchronization (ERD/ERS) patterns during kinesthetic motor imagery. (A) Grand-average mu ERD/ERS [%] time courses averaged across the four sensorimotor electrodes of interest (red electrodes shown in Panel B) for the MVF and sham group during MI1 and MI2. Shaded areas indicate 95% confidence intervals. The dark gray shaded area indicates the baseline interval (−1000 to −500 ms), and the light gray shaded area indicates the predefined analysis window (500–2500 ms). Vertical dashed lines indicate trial onset (0 ms) and the end of the imagery period (6000 ms). Negative values reflect ERD, and positive values reflect ERS. (B) Grand-average scalp topographies of mu ERD averaged across MI1 and MI2 and the predefined analysis window (500–2500 ms) relative to baseline. Negative values (blue) indicate ERD, and positive values (red) indicate ERS. Red dots indicate the four sensorimotor electrodes included in the primary analyses. (C) Grand-average time–frequency representations averaged across the same sensorimotor electrodes and across MI1 and MI2 for the MVF (left) and sham (right) group. The color scale represents percentage power change relative to baseline. Negative values (blue) indicate ERD, and positive values (red) indicate ERS. Vertical dashed lines indicate trial onset (0 ms) and the end of the imagery period (6000 ms). MVF = mirror visual feedback; MI = motor imagery.

#### 2.6.1. Independent Component Analysis and Bad Channel Detection

Before ICA, the three EEG datasets (MI1, Intervention, MI2) acquired from each participant were concatenated and subsequently low-pass filtered at 40 Hz (transition bandwidth = 10 Hz, order = 84) as well as high-pass filtered at 1 Hz (transition bandwidth = 2 Hz, order = 414) using Hamming-windowed FIR filters. Channels whose signal standard deviation exceeded the mean channel standard deviation by more than 2.5 standard deviations across channels were classified as bad and removed. The data were segmented into consecutive 1-s epochs, and epochs containing artifacts were excluded using EEGLAB’s pop_jointprob function (*SD* = 2). ICA decomposition was performed on the cleaned concatenated data using the extended infomax algorithm. The resulting unmixing weights were applied to the original, unfiltered and non-concatenated EEG datasets with the same bad channels removed. Independent components reflecting artefacts were identified using a combination of automated classification and visual inspection. Automated classification was performed using ICLabel (Pion-Tonachini et al., 2019) and Eye-Catch (Bigdely-Shamlo et al., 2013). ICLabel components classified as eye-related with a probability >70% or as muscle- or heart-related with a probability >80% were automatically rejected. Components identified as ocular artifacts by Eye-Catch were additionally excluded. ICs not flagged by ICLabel or Eye-Catch were visually inspected based on their scalp topographies, power spectra, and time courses. Components showing topographical patterns characteristic of ocular artefacts (e.g., strong frontal activity or bipolar frontal distributions; Villasana, 2022) were additionally rejected when their spectral and temporal characteristics were consistent with non-neural activity. On average, 2.55 ± 0.64 ICs (range: 2–5)^1^ were removed per dataset. Bad channels were subsequently interpolated using spherical interpolation. To ensure consistent linked-mastoid rereferencing, both mastoid channels were interpolated whenever one mastoid channel had been identified as bad. On average, 1.05 ± 0.90 channels (range: 0–2)^1^ were interpolated per participant.

#### 2.6.2. Preprocessing and Epoch Rejection

The cleaned continuous data were low-pass filtered at 40 Hz using a Hamming-windowed FIR filter (transition bandwidth = 10 Hz, order = 84) and high-pass filtered at 1 Hz using a Hamming-windowed FIR filter (transition bandwidth = 2 Hz, order = 414). After re-referencing to the average of the left and right mastoids, continuous EEG data were segmented into epochs from −2500 to 9000 ms relative to MI trial onset. Timepoint 0 corresponded to the onset of the instruction screen specifying the finger opposition movement and pronation/supination combination. Using EEGLAB’s pop_jointprob function (*SD* = 4), epochs containing residual artefacts were automatically flagged. The result was checked visually for missed artefacts at the channels of interest within the interval −1500 to 7000 ms, and if additional artefacts were detected, the corresponding epochs were similarly flagged. This information was aggregated with the information regarding epochs flagged by EMG analysis, and all flagged epochs were excluded. On average, 4.21 ± 4.56 epochs were excluded per dataset (range: 0-29 out of 60 trials). Following epoch rejection, participants retained an average of 55.78 ± 3.02 out of 60 trials (range: 49–60) for the ERD analysis in the MVF group during MI1, 55.39 ± 4.96 out of 60 trials (range: 41–60) during MI2, 55.35 ± 6.47 out of 60 trials (range: 31–60) in the sham group during MI1, and 56.60 ± 2.96 out of 60 trials (range: 49–60) during MI2.^1^

#### 2.6.3. Quantification of ERD/ERS

Time-frequency representations of the preprocessed EEG signal were obtained using the EEGLAB function pop_newtimef. Analyses focused on four central and centro-parietal sensorimotor electrodes (cf. section 2.6). For each MI dataset, time-resolved spectral power was obtained using a complex Morlet wavelet transform. Frequencies between 5 and 35 Hz were sampled logarithmically, with the number of cycles increasing linearly from 3 at the lowest frequency to 10 at the highest frequency. The analysis window spanned -2500 to 8996 ms. Baseline correction within EEGLAB was disabled because baseline normalization was applied manually within the ERD/ERS formula. Power estimates P(f,t,trial) were computed as the squared magnitude of the complex wavelet coefficients of the single-trial data. The trial-averaged baseline power R(f) was obtained by averaging power across trials within the pre-trial reference (baseline) interval. The baseline interval was defined as -1000 ms to -500 ms relative to the onset of the MI instruction cue. ERD/ERS values were then calculated for each trial separately using this trial-averaged baseline, applying an adaptation of the percentage change formula of Pfurtscheller and Lopes da Silva (1999): 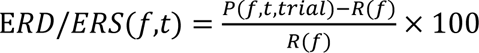. Because R(f) was constant across trials, this single-trial computation is mathematically equivalent to computing the original formula on the trial-averaged power. After normalization, ERD/ERS values were averaged across trials for each electrode and session. Mean ERD/ERS values were extracted from the mu band (8–12 Hz) and a time window from 500 to 2500 ms. The upper bound of the analysis window was determined from the grand-average mu ERD recorded at the left sensorimotor electrodes, computed across all available participants prior to ERD-based exclusions (*N* = 40). The window extended until the signal had recovered to 50% of its peak ERD, thereby capturing the main ERD period while minimizing the inclusion of later rebound effects.

### 2.7. Statistical Analysis

#### 2.7.1. Behavioral Data

MI difficulty ratings and their changes within sessions were analyzed using a 2 × 2 × 2 mixed-design ANOVA with group (MVF, sham) as the between-subject factor and MI session (MI1, MI2) and session half (first, second) as within-subject factors. Attention and tiredness ratings collected during the intervention were analyzed separately using two 2 × 3 mixed-design ANOVAs with group (MVF, sham) as the between-subject factor and intervention block (Block 1, Block 2, Block 3) as the within-subject factor. Likert scale items from the post-intervention and post-procedure questionnaires were analyzed using independent-samples t-tests. For items that were only completed by the MVF group, descriptive statistics are reported.

#### 2.7.2. EEG Data

To test whether the increase in mu ERD during kMI from pre- to post-intervention is greater in the MVF group than in the sham group (Hypothesis 1), mu-band ERD was analyzed using a 2 × 2 × 2 mixed-design ANOVA with MI session (MI1, MI2) and hemisphere (left, right) as within-subject factors and group (MVF, sham) as the between-subject factor. To derive the dependent variable for each hemisphere, mu ERD values from the predefined sensorimotor electrodes of interest were averaged within each hemisphere. To test whether the increase in mu ERD lateralization toward contralateral sensorimotor areas during kMI from pre- to post-intervention is greater in the MVF group than in the sham group (Hypothesis 2), a 2 × 2 mixed-design ANOVA was performed with MI session (MI1, MI2) as the within-subject factor and group (MVF, sham) as the between-subject factor. Mu ERD lateralization was calculated as the difference between left- and right hemisphere mu ERD values (LH – RH), based on hemisphere-averaged values derived from the electrodes of interest. All statistical analyses were conducted in JASP (JASP Team, 2025, version 0.95.4). Assumptions of normality were assessed using the Shapiro-Wilk test. Parametric analyses were used when assumptions were met. When assumptions were violated, the corresponding non-parametric alternative was applied where available. Additional exploratory analyses were conducted to further characterize potential effects of MVF beyond the predefined hypotheses. First, Bayesian analogues of the primary frequentist ANOVAs were performed to quantify evidence for the inclusion or exclusion of the hypothesized effects. Second, a cluster-based permutation analysis (Maris & Oostenveld, 2007) was conducted within the MVF group to examine pre- to post-intervention changes in oscillatory activity without restricting the analysis to the predefined channels, frequency range, or time window used in the primary analyses. The analysis was performed in FieldTrip using a dependent-samples t-test (ft_statfun_depsamplesT) and cluster-based correction for multiple comparisons. All non-mastoid EEG channels, frequencies from 5 to 35 Hz, and the time interval from −2500 to 7000 ms relative to the onset of the motor MI instruction cue were included. Clusters were formed across neighbouring electrodes, frequencies, and time points, with no minimum number of neighbouring channels required for cluster inclusion, and statistical significance was assessed using a Monte Carlo permutation procedure with 5000 random permutations. Third, exploratory correlation analyses were conducted to examine associations between subjective intervention ratings and changes from MI1 to MI2 in contralateral mu ERD as well as mu ERD lateralization. Correlations were computed within the MVF group and, for ratings available in both groups, also within the sham group. Correlation coefficients were accompanied by 95% bootstrap confidence intervals based on 5,000 bootstrap samples. Descriptive statistics are reported as mean ± *SD*. For inferential analyses, means are reported with their corresponding standard errors (*SE*). Ninety-five percent confidence intervals (95% CIs) are additionally reported where appropriate. Effect sizes are reported as partial eta squared (*η_p_*²) for ANOVAs and rank biserial correlation (*r*) for the Mann-Whitney U Test. The alpha level was set to .05 for all analyses.

## 3. Results

### 3.1. Behavioral Results

Behavioral ratings were generally comparable between the MVF and sham groups. Participants rated kMI as moderately easy in both MI sessions. During the intervention, ratings for attention to the intervention-specific visual input were high and tiredness ratings were low across groups and blocks. Participants in the MVF group generally reported a realistic mirror illusion. Post-experimental ratings further indicated that the task instructions were perceived as clear and that the overall procedure was moderately demanding.

The 2 × 2 × 2 mixed-design ANOVA of MI difficulty ratings revealed no significant main effects of group, MI session, or session half, and no significant interactions (all *p* > .05; Table 1). Attention and tiredness ratings remained stable across intervention blocks and did not differ between groups. Separate mixed-design ANOVAs revealed no significant main effects or interactions for either measure (all *p* > .05; Table 2). Questionnaire items assessing the subjective experience of the mirror illusion were administered only to the MVF group and were therefore analyzed descriptively. Descriptive statistics are reported in Table 3. For post-procedure questionnaire items completed by both groups, Mann-Whitney U tests revealed no significant group differences (all *p* > .05). Detailed descriptive statistics and complete statistical test results are provided in Supplementary Tables S1–S5.

**Table 1:** Mixed-Design ANOVA Results for MI Difficulty Ratings by MI Session and Session Half.

| Effect | df | *F* | *p* | *η<sub>p</sub>*<sup>2</sup> |
|---|---|---|---|---|
| MI Session | 1, 36 | 0.08 | .782 | 0.002 |
| Group | 1, 36 | 1.01 | .321 | 0.027 |
| Half | 1, 36 | 0.70 | .410 | 0.019 |
| MI Session × Group | 1, 36 | 2.47 | .125 | 0.064 |
| Half × Group | 1, 36 | 0.38 | .540 | 0.011 |
| MI Session × Half | 1, 36 | 3.03 | .090 | 0.078 |
| MI Session × Half × Group | 1, 36 | 0.44 | .514 | 0.012 |

MI difficulty rating was obtained on a 7-point Likert scale (1 = very hard, 7 = very easy).

**Table 2:** Mixed-Design ANOVA Results for Attention and Tiredness Ratings by Intervention Block.

| Effect | df | *F* | *p* | *η<sub>p</sub>*<sup>2</sup> |
|---|---|---|---|---|
| **Attention** |  |  |  |  |
| Block | 2, 72 | 0.75 | .476 | 0.020 |
| Group | 1, 36 | 0.83 | .368 | 0.023 |
| Block × Group | 2, 72 | 0.05 | .948 | 0.001 |
| **Tiredness** |  |  |  |  |
| Block | 2, 72 | 1.32 | .273 | 0.035 |
| Group | 1, 36 | 1.09 | .303 | 0.029 |
| Block × Group | 2, 72 | 0.67 | .514 | 0.018 |

Ratings for attention and tiredness were obtained on 7-point Likert scales (1 = strongly disagree, 7 = strongly agree).

**Table 3:**
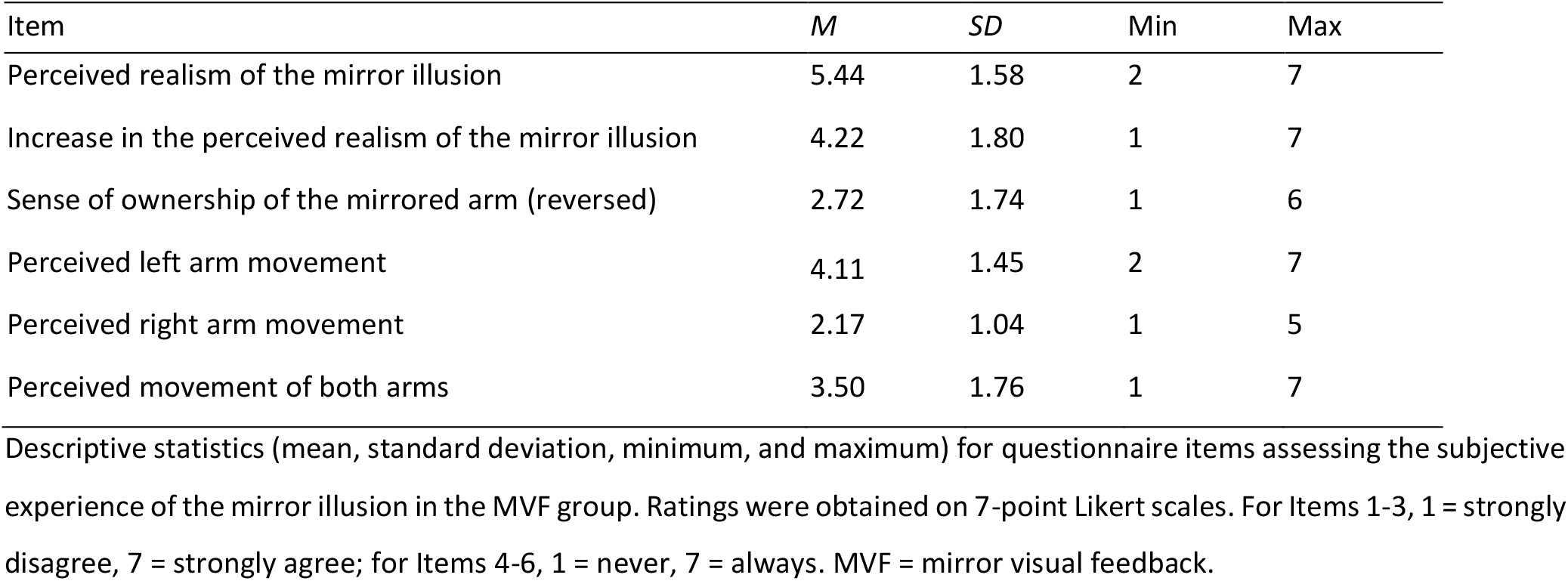
Descriptive statistics of MVF intervention questionnaire items.

| Item | *M* | *SD* | Min | Max |
|---|---|---|---|---|
| Perceived realism of the mirror illusion | 5.44 | 1.58 | 2 | 7 |
| Increase in the perceived realism of the mirror illusion | 4.22 | 1.80 | 1 | 7 |
| Sense of ownership of the mirrored arm (reversed) | 2.72 | 1.74 | 1 | 6 |
| Perceived left arm movement | 4.11 | 1.45 | 2 | 7 |
| Perceived right arm movement | 2.17 | 1.04 | 1 | 5 |
| Perceived movement of both arms | 3.50 | 1.76 | 1 | 7 |

Descriptive statistics (mean, standard deviation, minimum, and maximum) for questionnaire items assessing the subjective experience of the mirror illusion in the MVF group. Ratings were obtained on 7-point Likert scales. For Items 1-3, 1 = strongly disagree, 7 = strongly agree; for Items 4-6, 1 = never, 7 = always. MVF = mirror visual feedback.

### 3.2. Electrophysiological Results

The kMI task elicited the expected pattern of sensorimotor rhythm desynchronization during imagery and rebound synchronization following imagery termination. The grand-average time courses for all four group-by-session conditions (Figure 2A) showed a clear and sustained mu ERD during the imagery period. Mu power decreased shortly after MI trial onset, reached its maximum desynchronization within the predefined analysis window (500–2500 ms), and subsequently increased after imagery offset, reflecting a post-imagery ERS rebound. Visual inspection indicated highly similar temporal profiles across groups and sessions.

The scalp topographies averaged across the mu band (8–12 Hz) and analysis window (Figure 2B) revealed a pronounced left-lateralized distribution of mu ERD in both groups. Although maximal desynchronization was observed over centro-parietal regions, the effect extended into channels covering adjacent parietal areas.

The time-frequency representations averaged across the electrodes of interest (Figure 2C) further confirmed a distinct reduction in power within the mu frequency range during the imagery period, followed by a subsequent power increase after imagery offset. In addition, desynchronization extended into the lower beta range, and both groups showed a further localized reduction in power within the higher beta range.

To test whether the increase in mu ERD during kMI from pre- to post-intervention is greater in the MVF group than in the sham group (Hypothesis 1), mean mu ERD values were analyzed using a 2 × 2 × 2 mixed-design ANOVA. Figure 3A illustrates the distribution of mean mu ERD values across groups, sessions, and hemispheres, and a full overview of all frequentist effects is provided in Table 4. No significant main effects of MI session or group were observed, and none of the interaction effects reached significance (all *p* > .05). Thus, Hypothesis 1 was not supported. However, a significant main effect of hemisphere was found, *F*_(1, 36)_ = 28.85, *p* < .001, *η_p_*² = .445, with stronger mu ERD in the left hemisphere (*M* = -25.36 %, *SE* = 2.59) than in the right hemisphere (*M* = -16.22 %, *SE* = 3.06). The complementary Bayesian ANOVA yielded a BF_incl_ of 0.647 for the predicted MI session × group interaction, lending anecdotal evidence against inclusion of this interaction. By contrast, there was strong evidence for inclusion of the main effect of hemisphere (BF_incl_ = 5629.018). A full overview of the Bayesian results is provided in Supplementary Table S6.

**Figure 3:**
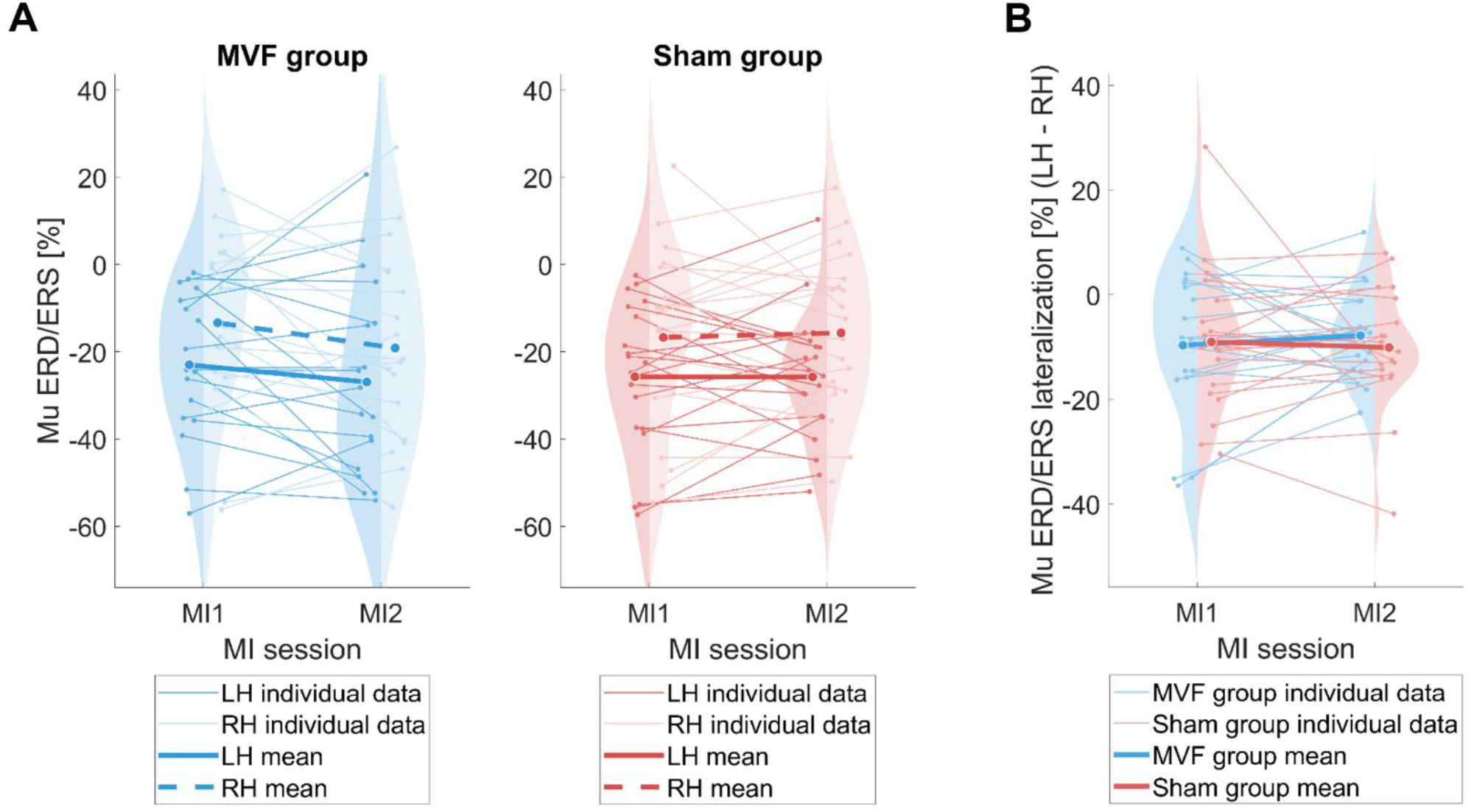
Distribution of mu event-related desynchronization (ERD) and mu ERD lateralization across MI sessions. (A) Distribution of mean mu ERD [%] during MI1 and MI2 for the MVF (left) and sham (right) group, shown separately for the left (LH) and right (RH) hemispheres. Negative values reflect ERD, whereas positive values reflect event-related synchronization (ERS). (B) Distribution of mu ERD lateralization scores [%], calculated as the difference between left- and right-hemispheric mu ERD values (LH − RH). Negative lateralization values indicate stronger mu ERD over the left hemisphere, whereas positive values indicate stronger mu ERD over the right hemisphere. Violin plots represent the distribution of participant values, thin lines indicate individual participants across sessions, and larger markers represent group means. Mu ERD values were averaged across the predefined analysis window (500–2500 ms) relative to baseline (−1000 to −500 ms) and the sensorimotor electrodes of interest within each hemisphere. MVF = mirror visual feedback; MI = motor imagery.

**Table 4:** Results of the frequentist mixed-design ANOVA on mu ERD.

| Effect | df | *F* | *p* | *η<sub>p</sub>*<sup>2</sup> |
|---|---|---|---|---|
| MI Session | 1, 36 | 0.88 | .354 | 0.024 |
| Group | 1, 36 | 0.01 | .946 | 1.285x10<sup>-4</sup> |
| Hemisphere | 1, 36 | 28.85 | < .001 | 0.445 |
| MI Session × Group | 1, 36 | 1.34 | .255 | 0.036 |
| Hemisphere × Group | 1, 36 | 0.06 | .806 | 0.002 |
| MI Session × Hemisphere | 1, 36 | 0.04 | .844 | 0.001 |
| MI Session × Hemisphere × Group | 1, 36 | 0.54 | .469 | 0.015 |

Results of a 2 × 2 × 2 frequentist mixed-design ANOVA with MI session and hemisphere as within-subject factors and group as the between-subject factor.

To test whether the increase in mu ERD lateralization toward contralateral sensorimotor areas during kMI from pre- to post-intervention is greater in the MVF group than in the sham group (Hypothesis 2), mu ERD lateralization scores were analyzed using a 2 × 2 mixed-design ANOVA. Neither the main effect of MI session nor the main effect of group reached significance. Crucially, the MI session × group interaction was also not significant (all *p* > .05; Table 5, Figure 3B). Therefore, Hypothesis 2 was not supported. The complementary Bayesian ANOVA yielded a BF_incl_ of 0.386 for the MI session × group interaction, indicating anecdotal evidence against inclusion of the interaction. A full overview of the Bayesian results is provided in Supplementary Table S7.

**Table 5:**
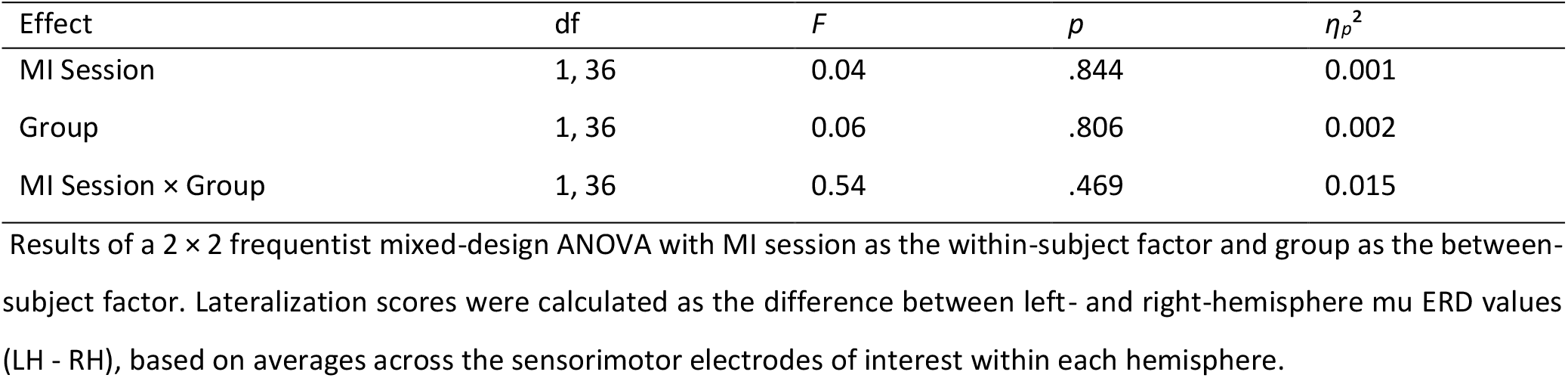
Results of the frequentist mixed-design ANOVA on mu ERD lateralization.

| Effect | df | *F* | *p* | *η<sub>p</sub>*<sup>2</sup> |
|---|---|---|---|---|
| MI Session | 1, 36 | 0.04 | .844 | 0.001 |
| Group | 1, 36 | 0.06 | .806 | 0.002 |
| MI Session × Group | 1, 36 | 0.54 | .469 | 0.015 |

Results of a 2 × 2 frequentist mixed-design ANOVA with MI session as the within-subject factor and group as the between-subject factor. Lateralization scores were calculated as the difference between left- and right-hemisphere mu ERD values (LH - RH), based on averages across the sensorimotor electrodes of interest within each hemisphere.

### 3.3. Exploratory Analyses

To explore whether MVF-related changes occurred outside the predefined analysis window, frequency range, and channels, a cluster-based permutation analysis was conducted. No significant positive or negative clusters were identified after correction for multiple comparisons (5000 permutations; all *p* > .05). Thus, within the MVF group, no spatially, spectrally, and temporally contiguous patterns of oscillatory activity significantly differed between MI1 and MI2.

Exploratory Spearman correlations were computed between subjective intervention ratings and changes from MI1 to MI2 in contralateral mu ERD and mu ERD lateralization (Figure 4). The ratings reported below were assessed on 7-point scales ranging from 1 (strongly disagree) to 7 (strongly agree), with higher values reflecting greater illusion realism, tiredness, or attention to the intervention-specific visual input, respectively. Overall, associations varied in magnitude, ranging from negligible to moderate. For contralateral mu ERD, the largest association in the MVF group was observed between illusion realism and mu ERD change (*ρ* = .44, 95% bootstrap CI [−.06, .82], *p* = .066), with participants reporting a more realistic mirror illusion tending to show a weaker contralateral mu ERD during MI2 relative to MI1. In the sham group, the largest association was observed between tiredness and mu ERD change (*ρ* = −.42, 95% bootstrap CI [−.71, .02], *p* = .063), with participants reporting greater tiredness tending to show a stronger contralateral mu ERD during MI2 relative to MI1. For mu ERD lateralization, the most notable pattern concerned attention, which showed associations in opposite directions in the MVF (*ρ* = −.45, 95% bootstrap CI [−.82, .07], *p* = .059) and sham group (*ρ* = .43, 95% bootstrap CI [.06, .67], *p* = .061). Greater attention to the mirrored upper extremity in the MVF group was associated with a greater shift in lateralization toward the contralateral hemisphere from MI1 to MI2, whereas greater attention to the wooden panel in the sham group was associated with a smaller shift toward the contralateral hemisphere. Figure 4 provides a visual overview of all correlations, and Supplementary Tables S8 and S9 report the corresponding *ρ* values, 95% bootstrap confidence intervals, and *p* values.

**Figure 4:**
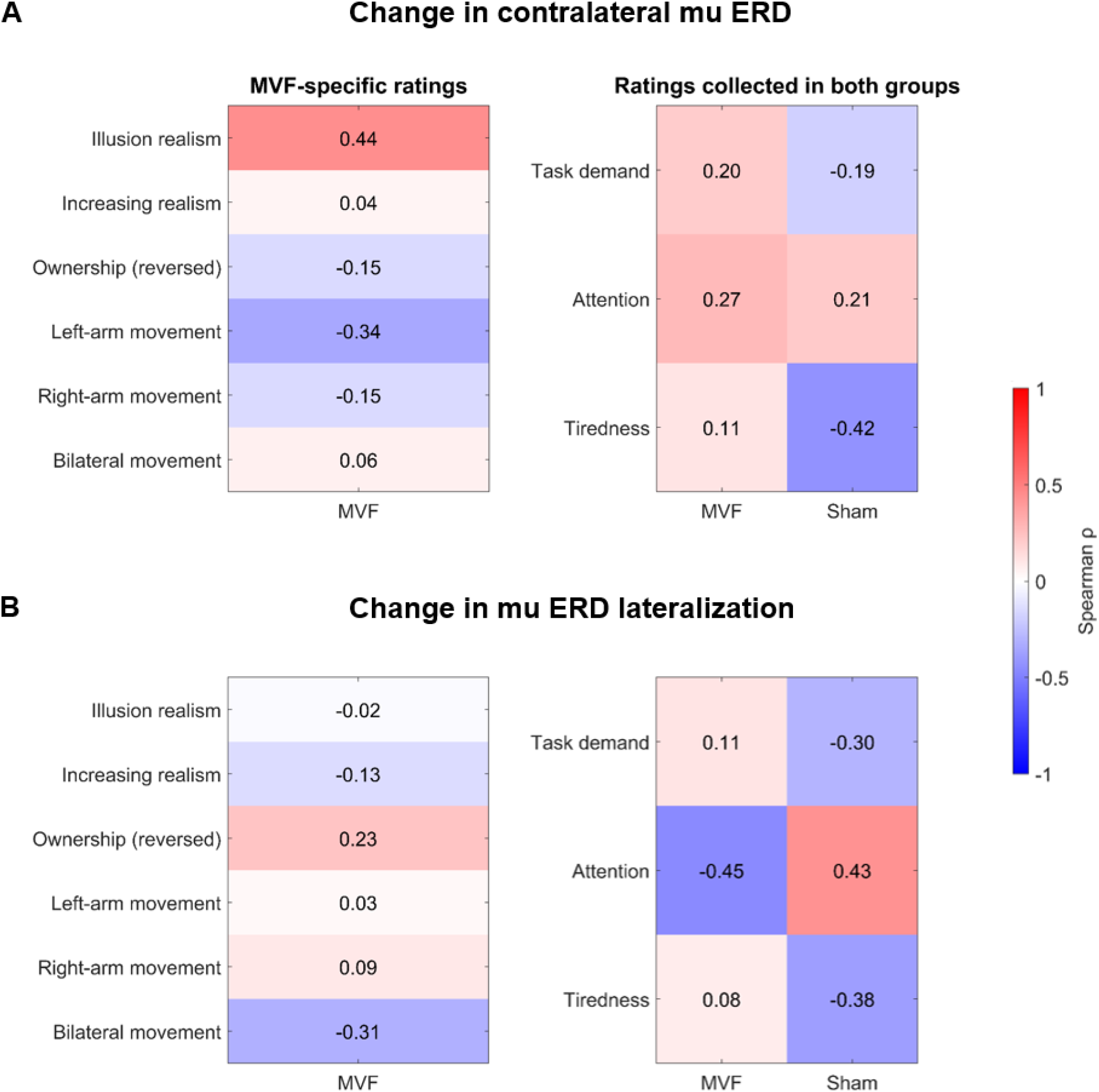
Spearman correlations between subjective intervention ratings and changes from MI1 to MI2 in (A) contralateral mu event-related desynchronization (ERD) and (B) mu ERD lateralization. Left panels show correlations with MVF-specific ratings collected only in the MVF group, whereas right panels show correlations with ratings collected in both the MVF and sham groups. Cell colors represent Spearman’s correlation coefficient (*ρ*), with red indicating positive correlations and blue indicating negative correlations. (A) Positive coefficients indicate that higher questionnaire ratings were associated with larger (less negative) mu ERD change scores, whereas negative coefficients indicate associations with smaller (more negative) change scores, reflecting stronger contralateral mu ERD during MI2 relative to MI1. (B) Positive coefficients indicate that higher questionnaire ratings were associated with larger (less negative) mu ERD lateralization change scores, whereas negative coefficients indicate associations with smaller (more negative) change scores, reflecting a shift in lateralization toward the contralateral left hemisphere from MI1 to MI2. Ratings were obtained on 7-point Likert scales. For illusion realism, increasing realism, ownership, task demand, attention, and tiredness, ratings ranged from 1 = strongly disagree to 7 = strongly agree. For left-arm movement, right-arm movement, and bilateral movement, ratings ranged from 1 = never to 7 = always. The attention item referred to attentive observation of the arm in the mirror in the MVF group and of the wooden panel in the sham group. The ownership item was negatively worded. Thus, higher scores indicate lower perceived ownership. MVF = mirror visual feedback.

## 4. Discussion

The present study investigated whether a single session of MVF modulates the neural correlates of subsequent kMI, as reflected by sensorimotor mu ERD in healthy young adults. Based on previous evidence suggesting functional overlap between MVF and MI, and synergistic effects when both processes are combined (Bello et al., 2020; Fukumura et al., 2007; Gómez & Braidot, 2024; Kang et al., 2011), our predictions were twofold. First, we hypothesized that the increase in mu ERD from pre- to post-intervention would be greater in the MVF group than in the sham group. Second, the MVF group was expected to show a greater increase in mu ERD lateralization toward contralateral sensorimotor areas during kMI compared to the sham group.

Across groups and sessions, the kMI task reliably elicited the expected mu ERD over sensorimotor regions, suggesting successful engagement of sensorimotor networks. In addition, mu ERD was stronger over the contralateral left than the right hemisphere. Contrary to our hypotheses, however, a single session of MVF did not enhance subsequent kMI-related mu ERD or its lateralization relative to the sham intervention. Thus, the present findings do not support the assumption that a brief MVF intervention modulates the neural correlates of subsequent kMI.

### 4.1. Absence of MVF-related Modulation of Subsequent kMI

Our findings do not support that a single session of MVF modulates mu ERD during subsequent kMI. Previous studies have reported synergistic effects for corticospinal excitability when MI and MVF are combined (Fukumura et al., 2007; Kang et al., 2011). The absence of an effect of MVF on MI-related neural activity in the present study may appear to contrast with these findings, but this apparent discrepancy may be explained by important methodological differences. Specifically, these studies examined MI performed concurrently with MVF and therefore cannot determine whether MVF induces lasting changes that influence subsequent MI. The facilitation observed in these paradigms may depend on the concurrent integration of visual information with MI rather than on a sustained priming of sensorimotor areas induced by a single session of MVF.

Gómez and Braidot (2024) more directly addressed the possibility of offline MVF effects on subsequent MI. Using an *r*²-based spectral measure that quantified how strongly EEG activity during MI differed from the control condition, they reported greater MI-related modulation when MVF preceded rather than followed MI. However, several methodological aspects of their design complicate the interpretation of these findings. First, the study employed a between-group design, comparing participants who performed MVF before MI with those who performed MVF after MI. Second, the groups differed not only in the timing of MVF exposure but also in the amount of motor execution performed prior to MI. Participants in the MVF-first group completed an additional block of movement practice before the imagery task. Consequently, rather than MVF-specific mechanisms, the reported enhancement of MI-related activity may reflect effects of prior motor execution, task familiarity, or sensorimotor priming. Such alternative interpretation would be consistent with studies showing facilitative effects of physical practice on subsequent MI-related neural or behavioral measures (Williams et al., 2011; Wriessnegger et al., 2018; Wriessnegger et al., 2014). Taken together, the discrepancy between Gómez and Braidot (2024) and the present findings may reflect differences in experimental design rather than genuine evidence for an offline MVF effect on subsequent MI. By controlling for baseline differences and prior motor execution, the present study provides a more stringent test of this hypothesis.

The absence of the predicted effects may reflect a genuine lack of offline MVF modulation of subsequent kMI, although the Bayesian analyses provide only limited support for this interpretation. The Bayesian ANOVAs yielded anecdotal evidence against the predicted MI Session × Group interactions for both mu ERD and mu ERD lateralization. Thus, while the findings do not support the hypothesized MVF-related effects, they also do not provide strong evidence for their absence. One possible explanation for the lack of an observable offline effect concerns the modality-specific characteristics of MVF and kMI. MVF primarily relies on visual feedback and the induction of a visual illusion (Deconinck et al., 2015; Thieme et al., 2018), whereas kMI is based on internally generated movement sensations (Dickstein & Deutsch, 2007). This mismatch between externally driven visual input and internally generated kinesthetic representations may therefore have limited transfer effects. Notably, as Gómez and Braidot (2024) did not specify which imagery modality their participants used during the MI condition, it remains unclear whether this mismatch also applied to the reported offline effect of MVF on MI. In the present study, kMI rather than vMI was deliberately used, as it is known to more strongly engage sensorimotor areas that are also involved in MVF (Bello et al., 2020). Moreover, kMI has been suggested to yield greater benefits in motor rehabilitation contexts than vMI (Kim et al., 2011). Thus, kMI appeared to provide a theoretically promising basis for potential transfer effects while also aligning with the broader goal of optimizing stroke rehabilitation. Nevertheless, despite these considerations, the difference in dominant sensory modality may have limited the transfer of MVF-related processes to subsequent kMI.

A further potential explanation for the null findings concerns the temporal characteristics of MVF interventions. The Berlin Mirror Therapy Protocol (Morkisch & Dohle, 2015) recommends repeated training over multiple sessions (e.g., around 20 sessions). Similarly, a systematic review by Thieme and colleagues (2018) included studies in which mirror therapy was administered between three and seven days per week over periods of two to eight weeks, with session durations ranging from 15 to 60 min. This suggests that MVF may require repeated application before influencing subsequent MI. In contrast, the present study aimed to detect immediate effects of a single MVF session on subsequent kMI.

From a conceptual perspective, it is important to note that an increase in mu ERD does not necessarily reflect improvements in MI performance. While mu ERD is commonly interpreted as a correlate of increased cortical activation (Klimesch, 2012), its relationship to task performance is not strictly linear. In the context of motor learning, increasing expertise and automatization are often associated with reduced cortical activation despite improved behavioral performance, a phenomenon commonly referred to as neural efficiency (Picard et al., 2013; Wright et al., 2012). In line with this, a study on MI has shown that superior imagery performance can be accompanied by decreased activation in motor-related cortical areas, suggesting more efficient internal simulation processes (Zhang et al., 2019). Accordingly, the absence of increased ERD following MVF does not necessarily imply the absence of beneficial effects on MI-related processing, but may instead reflect stable or more efficient neural processing.

Overall, the absence of MVF effects in the present study may reflect a combination of modality-specific differences, insufficient exposure duration, and the complex relationship between neural activation and imagery performance. However, it is also important to consider that the present findings may reflect a genuine absence of offline modulation of MI-related mu ERD by MVF. While several theoretical and empirical considerations suggest a potential interaction between MVF and MI (Bello et al., 2020; Deconinck et al., 2015; Fukumura et al., 2007; Gómez & Braidot, 2024; Kang et al., 2011), the present results do not support the assumption that a brief MVF intervention systematically alters subsequent MI-related neural activity.

### 4.2. Additional Evidence from Exploratory Analyses

The exploratory analyses were conducted to further evaluate the robustness of the primary findings and to examine whether changes in MI-related neural activity from MI1 to MI2 were associated with subjective responses to the intervention.

With respect to channel selection, the electrodes included in the primary analyses were defined a priori based on previous studies investigating MI in combination with MVF or visual input (Gómez & Braidot, 2024; M. Lin et al., 2023; Ono et al., 2013). However, the scalp topographies in Figure 2B showed that mu ERD was predominantly distributed over more postcentral regions. This spatial distribution is consistent with previous findings showing mu ERD centered over centro-parietal electrodes (Daeglau et al., 2020) and MEG evidence localizing mu activity to postcentral cortical regions (Cheyne, 2013). However, because the cluster-based permutation analysis was not restricted to the predefined ROI and revealed no significant clusters across the analyzed electrodes, the results suggest that using a different, more postcentrally located ROI would not have altered the conclusion regarding the absence of an MVF-related effect.

With respect to frequency, the cluster-based permutation analysis revealed no significant clusters across the analyzed frequency range (5-35 Hz). This indicates that potential MVF-related effects were not present in frequency bands outside the predefined mu range. Within the mu frequency range, ERD is commonly regarded as a neural correlate of sensorimotor processing and it can be subdivided into functionally distinct low-mu (8–10 Hz) and high-mu (10–12 Hz) bands (Hervault et al., 2021; Pfurtscheller et al., 2000). Lower mu ERD has been described as being more spatially widespread and less somatotopically specific, whereas higher mu ERD is more spatially focal and somatotopically specific (Pfurtscheller et al., 2000). In addition, many MI studies, including Gómez and Braidot (2024), analyze the broader sensorimotor rhythm, encompassing both mu and beta frequencies. Beta-band ERD has been linked more closely to motor preparation and control, and MEG findings indicate that beta activity is primarily localized to central cortical regions (Cheyne, 2013; Hari & Salmelin, 1997).

With respect to time, the cluster-based permutation analysis revealed no significant clusters throughout the analyzed epoch. The primary analysis window was selected in a data-driven manner based on the grand-average, cross-condition, mu ERD time course for the left sensorimotor electrodes. The analysis window was then tailored to the ERD time course and values were averaged over time. This approach might have obscured short-lived mu effects. The exploratory results suggest this to be unlikely. Likewise, effects occurring in different frequency bands or spatial locations might have been characterized by different temporal dynamics. The absence of significant clusters across the analyzed epoch also argues against these possibilities.

Whereas the cluster-based permutation analysis addressed the robustness of the primary findings, the exploratory correlation analyses examined whether individual differences in the subjective experience of the MVF intervention or more general time on task effects were associated with changes in MI-related neural activity. Interestingly, the strongest correlation regarding the change in mu ERD was observed for illusion realism, with participants reporting a more realistic mirror illusion tending to show a weaker contralateral mu ERD during MI2 relative to MI1. Although this pattern does not support the original hypothesis, it is broadly compatible with the neural efficiency hypothesis (Picard et al., 2013; Wright et al., 2012), according to which greater task proficiency may be associated with reduced cortical recruitment. One speculative interpretation is that participants who were more susceptible to the mirror illusion may have engaged in subsequent imagery more efficiently, thereby requiring less sensorimotor recruitment during MI2.

For mu ERD lateralization, the most notable exploratory pattern concerned attention to the intervention-specific visual input. Greater attention to the mirrored upper extremity in the MVF group was associated with a greater shift in lateralization toward the contralateral hemisphere from MI1 to MI2, whereas greater attention to the wooden panel in the sham group showed the opposite association. These opposing patterns may point to a role of individual differences in attention to the intervention-specific visual input in subsequent sensorimotor lateralization.

Taken together, the exploratory analyses provide no indication that the primary null findings were attributable to the predefined spatial, spectral, or temporal analysis parameters, while the correlation analyses point to potential interindividual variability in the relationship between subjective intervention experiences and subsequent MI-related neural changes. Given their exploratory nature, however, these associations should be interpreted cautiously and require further investigation.

### 4.3. Towards Translational Stroke Rehabilitation Research

The present study was conducted in healthy participants, but its long-term motivation is to contribute to the optimization of stroke rehabilitation approaches. Both MVF and MI can engage sensorimotor networks without requiring movement of the corresponding limb (Deconinck et al., 2015; Munzert et al., 2009), making them particularly relevant for patients with substantial motor impairments who have limited capacity for physical practice. However, whether promoting ipsilesional recruitment is beneficial may depend on the individual pattern of post-stroke reorganization. The bimodal balance-recovery model proposes that the functional role of the contralesional hemisphere depends on the amount of preserved ipsilesional structural reserve (Di Pino et al., 2014). Whereas excessive interhemispheric inhibition from the contralesional hemisphere may impede recovery when structural reserve is relatively high, contralesional motor regions may assume a compensatory role when structural reserve is low. Consistent with this framework, Y.-L. Lin and colleagues (2020) demonstrated that the relationship between interhemispheric inhibition and motor impairment differs according to impairment severity. Accordingly, the neural and potentially therapeutic effects of MVF and MI may differ depending on patients’ structural reserve and impairment severity. Patients with sufficient ipsilesional structural reserve may particularly benefit from interventions supporting recruitment of the affected hemisphere, whereas in patients with extensive damage and low structural reserve, recovery may rely more strongly on compensatory contralesional mechanisms. This framework highlights that future studies investigating combined MVF and MI interventions should consider patient stratification based on ipsilesional structural reserve and impairment severity rather than assuming uniform treatment effects across stroke populations.

Beyond patient selection, successful translation also requires that experimental paradigms are feasible within the clinical setting. We conducted a feasibility testing of the present study protocol in three individuals in the chronic phase after stroke, all of which were familiar with MVF as implemented in the Berlin Mirror Therapy Protocol (Morkisch & Dohle, 2015). Despite extending the kMI period and dividing the task into shorter, more numerous blocks, none of the participants completed the protocol as originally planned. The main reason was not difficulties performing MI or engaging with the MVF intervention, but rather the overall session duration of 1:30 to 2:50 hours. While this may seem little surprising, it highlights a major challenge faced by experimental neurophysiology studies in clinical populations: reducing participant burden to a manageable amount, while maintaining a sufficient number of trials and data for reliable analyses (Stagneth et al., 2026).

### 4.4. Limitations

Several limitations should be considered when interpreting the results of the present study. These limitations concern the analytical approach and methodological aspects of the EEG recording.

With regard to the analysis, two limitations may have affected the sensitivity to detect MVF-related effects. First, in the present study, all artifact-free trials were included regardless of the presence or strength of a mu ERD. While this approach maximizes data inclusion, it may also introduce additional noise by including trials in which participants did not fully engage in the imagery task. Second, a fixed time window was used for ERD extraction. Although the cluster-based permutation analysis suggested that the primary findings were robust with respect to the analyzed time period, a fixed window may not fully capture individual variability in the onset and duration of MI across participants and trials. This variability may have been particularly pronounced in the present design, as participants were required to read and process task instructions before initiating imagery. Previous studies have addressed these issues by applying more fine-grained, trial-level analyses that account for variability in individual ERD time courses and trial characteristics (Daeglau et al., 2020). Such approaches may increase sensitivity to detect subtle MVF-related effects by reducing the influence of within- and between-subject variability. A further methodological consideration concerns the displacement of the EEG cap. Although the cluster-based permutation analysis suggests that the cap displacement did not materially affect the findings, some reduction in anatomical precision cannot be entirely excluded. Taken together, addressing these methodological limitations may further improve the sensitivity of future studies investigating the interaction between MVF and MI.

## 5. Conclusion

The present study investigated whether MVF modulates subsequent kMI-related neural activity. While kMI reliably elicited mu ERD, with stronger desynchronization over the contralateral hemisphere, no evidence was found that a single session of MVF enhanced subsequent MI, neither in terms of mu ERD nor its lateralization. These findings suggest that previously reported facilitatory effects of MVF on MI may depend on concurrent application and could, at least in part, reflect design choices rather than immediate aftereffects. At the same time, potential effects of repeated MVF exposure remain to be investigated. For future translational research, both patient stratification according to structural reserve and the development of feasible study designs tailored to severely impaired stroke populations should be considered. The present study provides a basis for future research aiming to optimize the combined use of MI and MVF in upper limb neurorehabilitation after stroke.

## Statements and Declarations

### Data and Code Availability Statement

The data that support the findings of this study are available from the corresponding author upon reasonable request. The MATLAB code is available on GitHub (https://github.com/linnschwarz/MI-MVF/tree/main).

### Ethics Approval Statement

The studies involving humans were approved by the Ethics Committee, Carl von Ossietzky Universität Oldenburg. The studies were conducted in accordance with the local legislation and institutional requirements. The participants provided their written informed consent to participate in this study.

### Permission to Reproduce Material from Other Sources

Permission to reproduce and adapt Figures from Appendix 1, p. 37 of Morkisch, N., & Dohle, C. (2015). *BeST - Berliner Spiegeltherapieprotokoll: Ein wissenschaftlich evaluiertes Manual zur Durchführung der Spiegeltherapie.* Hippocampus Verlag. https://www.hippocampus.de/Best.60317.html?shl=1 has been obtained from Hippocampus Verlag. The figure is reproduced and adapted with permission.

### CRediT Statement

LJS: Project administration, Conceptualization, Methodology, Investigation, Data curation, Formal analysis, Visualization, Software, Writing – original draft, Writing – review & editing. AS: Conceptualization, Project administration, Methodology, Resources, Supervision, Writing – review & editing. CK: Conceptualization, Funding acquisition, Project administration, Methodology, Resources, Supervision, Writing – review & editing.

## Acknowledgments

We thank Aaron Lohmann and Anja Stierl for their help in data acquisition. We also thank the center for Post-Acute Neurorehabilitation (P.A.N.) in Berlin for the opportunity to conduct exploratory feasibility sessions with individuals after stroke there as well as Dr. Dohle for enabling the data collection at the P.A.N. center and for his valuable feedback during the project.

Use of AI Generated Content (AIGC) and tools: During the preparation of this manuscript, the authors used ChatGPT (OpenAI, accessed October 2025 to July 2026) to assist with language editing, text refinement, code debugging, the implementation of selected analysis steps in scripts, and the generation of a methodological figure (Supplementary Figure S1). DeepL Write (DeepL GmbH) was used to improve grammar and wording. All AI-generated outputs were critically reviewed, verified, and revised where necessary. The authors take full responsibility for the content of this publication.

## Supplementary Material

The Supplementary Material for this article can be found online.

## Footnotes

1 Data for the final sample.

## References

ANT Neuro. (2025). Xensor digitizer software (Version 5.0.0) [Computer software].

Bai, Z., Fong, K. N. K., Zhang, J., & Hu, Z. (2020). Cortical mapping of mirror visual feedback training for unilateral upper extremity: A functional near-infrared spectroscopy study. Brain and Behavior, 10(1), e01489. 10.1002/brb3.1489

Bartur, G., Pratt, H., Dickstein, R., Frenkel-Toledo, S., Geva, A., & Soroker, N. (2015). Electrophysiological manifestations of mirror visual feedback during manual movement. Brain Research, 1606, 113–124. 10.1016/j.brainres.2015.02.029

Bartur, G., Pratt, H., Frenkel-Toledo, S., & Soroker, N. (2018). Neurophysiological effects of mirror visual feedback in stroke patients with unilateral hemispheric damage. Brain Research, 1700, 170–180. 10.1016/j.brainres.2018.09.003

Bello, U. M., Winser, S. J., & Chan, C. C. H. (2020). Role of kinaesthetic motor imagery in mirror-induced visual illusion as intervention in post-stroke rehabilitation. Reviews in the Neurosciences, 31(6), 659–674. 10.1515/revneuro-2019-0106

Bigdely-Shamlo, N., Kreutz-Delgado, K., Kothe, C., & Makeig, S. (2013). Eyecatch: Data-mining over half a million EEG independent components to construct a fully-automated eye-component detector. Annual International Conference of the IEEE Engineering in Medicine and Biology Society. IEEE Engineering in Medicine and Biology Society. Annual International Conference, 2013, 5845–5848. 10.1109/EMBC.2013.6610881

Chang, W. K., Lim, H., Park, S. H., Lim, C., Paik, N.-J., Kim, W.-S., & Ku, J. (2023). Effect of immersive virtual mirror visual feedback on Mu suppression and coherence in motor and parietal cortex in stroke. Scientific Reports, 13(1), 12514. 10.1038/s41598-023-38749-8

Cheyne, D. O. (2013). Meg studies of sensorimotor rhythms: A review. Experimental Neurology, 245, 27–39. 10.1016/j.expneurol.2012.08.030

Daeglau, M., Zich, C., Emkes, R., Welzel, J., Debener, S., & Kranczioch, C. (2020). Investigating Priming Effects of Physical Practice on Motor Imagery-Induced Event-Related Desynchronization. Frontiers in Psychology, 11, 57. 10.3389/fpsyg.2020.00057

Deconinck, F. J. A., Smorenburg, A. R. P., Benham, A., Ledebt, A., Feltham, M. G., & Savelsbergh, G. J. P. (2015). Reflections on mirror therapy: A systematic review of the effect of mirror visual feedback on the brain. Neurorehabilitation and Neural Repair, 29(4), 349–361. 10.1177/1545968314546134

Delorme, A., & Makeig, S. (2004). Eeglab: An open source toolbox for analysis of single-trial EEG dynamics including independent component analysis. Journal of Neuroscience Methods, 134(1), 9–21. 10.1016/j.jneumeth.2003.10.009

Di Pino, G., Pellegrino, G., Assenza, G., Capone, F., Ferreri, F., Formica, D., Ranieri, F., Tombini, M., Ziemann, U., Rothwell, J. C., & Di Lazzaro, V. (2014). Modulation of brain plasticity in stroke: A novel model for neurorehabilitation. Nature Reviews. Neurology, 10(10), 597–608. 10.1038/nrneurol.2014.162

Dickstein, R., & Deutsch, J. E. (2007). Motor imagery in physical therapist practice. Physical Therapy, 87(7), 942–953. 10.2522/ptj.20060331

Diers, M., Christmann, C., Koeppe, C., Ruf, M., & Flor, H. (2010). Mirrored, imagined and executed movements differentially activate sensorimotor cortex in amputees with and without phantom limb pain. Pain, 149(2), 296–304. 10.1016/j.pain.2010.02.020

Diers, M., Kamping, S., Kirsch, P., Rance, M., Bekrater-Bodmann, R., Foell, J., Trojan, J., Fuchs, X., Bach, F., Maaß, H., Cakmak, H., & Flor, H. (2015). Illusion-related brain activations: A new virtual reality mirror box system for use during functional magnetic resonance imaging. Brain Research, 1594, 173–182. 10.1016/j.brainres.2014.11.001

Fritzsch, C., Wang, J., Dos Santos, L. F., Mauritz, K.-H., Brunetti, M., & Dohle, C. (2014). Different effects of the mirror illusion on motor and somatosensory processing. Restorative Neurology and Neuroscience, 32(2), 269–280. 10.3233/RNN-130343

Fukumura, K., Sugawara, K., Tanabe, S., Ushiba, J., & Tomita, Y. (2007). Influence of mirror therapy on human motor cortex. The International Journal of Neuroscience, 117(7), 1039–1048. 10.1080/00207450600936841

Gómez, D. M. C., & Braidot, A. A. A. (2024). Improving motor imagery through a mirror box for BCI users. Journal of Neurophysiology, 131(5), 832–841. 10.1152/jn.00121.2023

Hari, R., & Salmelin, R. (1997). Human cortical oscillations: A neuromagnetic view through the skull. Trends in Neurosciences, 20(1), 44–49. 10.1016/S0166-2236(96)10065-5

Hervault, M., Zanone, P.-G., Buisson, J.-C., & Huys, R. (2021). Cortical sensorimotor activity in the execution and suppression of discrete and rhythmic movements. Scientific Reports, 11(1), 22364. 10.1038/s41598-021-01368-2

Ietswaart, M., Johnston, M., Dijkerman, H. C., Joice, S., Scott, C. L., MacWalter, R. S., & Hamilton, S. J. C. (2011). Mental practice with motor imagery in stroke recovery: Randomized controlled trial of efficacy. Brain: A Journal of Neurology, 134(Pt 5), 1373–1386. 10.1093/brain/awr077

ISWP. (2023). National Clinical Guideline for Stroke for the UK and Ireland. https://www.strokeguideline.org/contents/?_gl=1*1wo6b3e*_up*MQ.*_ga*MjA0OTc3NjE5OS4xNzc1NTAwMDQ2*_ga_EE3BZMVLRT*czE3NzU1MDAwNDUkbzEkZzAkdDE3NzU1MDAwNDUkajYwJGwwJGgw

JASP Team. (2025). *JASP (Version 0.95.4)* [Computer software]. https://jasp-stats.org

Jeannerod, M. (1995). Mental imagery in the motor context. Neuropsychologia, 33(11), 1419–1432. 10.1016/0028-3932(95)00073-c

Jeannerod, M. (2001). Neural simulation of action: A unifying mechanism for motor cognition. NeuroImage, 14(1 Pt 2), S103–9. 10.1006/nimg.2001.0832

Kang, Y. J., Ku, J., Kim, H. J., & Park, H. K. (2011). Facilitation of corticospinal excitability according to motor imagery and mirror therapy in healthy subjects and stroke patients. Annals of Rehabilitation Medicine, 35(6), 747–758. 10.5535/arm.2011.35.6.747

Kim, J.-S., Oh, D.-W., Kim, S.-Y., & Choi, J.-D. (2011). Visual and kinesthetic locomotor imagery training integrated with auditory step rhythm for walking performance of patients with chronic stroke. Clinical Rehabilitation, 25(2), 134–145. 10.1177/0269215510380822

Klimesch, W. (2012). Α-band oscillations, attention, and controlled access to stored information. Trends in Cognitive Sciences, 16(12), 606–617. 10.1016/j.tics.2012.10.007

Kothe, C., Shirazi, S. Y., Stenner, T., Medine, D., Boulay, C., Grivich, M. I., Artoni, F., Mullen, T., Delorme, A., & Makeig, S. (2025). The lab streaming layer for synchronized multimodal recording. Imaging Neuroscience (Cambridge, Mass.), 3. 10.1162/IMAG.a.136

Kwakkel, G., Stinear, C., Essers, B., Munoz-Novoa, M., Branscheidt, M., Cabanas-Valdés, R., Lakičević, S., Lampropoulou, S., Luft, A. R., Marque, P., Moore, S. A., Solomon, J. M., Swinnen, E., Turolla, A., Alt Murphy, M., & Verheyden, G. (2023). Motor rehabilitation after stroke: European Stroke Organisation (ESO) consensus-based definition and guiding framework. European Stroke Journal, 8(4), 880–894. 10.1177/23969873231191304

Langhorne, P., Coupar, F., & Pollock, A. (2009). Motor recovery after stroke: A systematic review. The Lancet. Neurology, 8(8), 741–754. 10.1016/S1474-4422(09)70150-4

Lee, H.-M., Li, P.-C., & Fan, S.-C. (2015). Delayed mirror visual feedback presented using a novel mirror therapy system enhances cortical activation in healthy adults. Journal of Neuroengineering and Rehabilitation, 12, 56. 10.1186/s12984-015-0053-1

Lim, V. K., Polych, M. A., Holländer, A., Byblow, W. D., Kirk, I. J., & Hamm, J. P. (2006). Kinesthetic but not visual imagery assists in normalizing the CNV in Parkinson’s disease. Clinical Neurophysiology: Official Journal of the International Federation of Clinical Neurophysiology, 117(10), 2308–2314. 10.1016/j.clinph.2006.06.713

Lin, M., Huang, J., Fu, J., Sun, Y., & Fang, Q [Qiang] (2023). A VR-Based Motor Imagery Training System With EMG-Based Real-Time Feedback for Post-Stroke Rehabilitation. IEEE Transactions on Neural Systems and Rehabilitation Engineering: A Publication of the IEEE Engineering in Medicine and Biology Society, 31, 1–10. 10.1109/TNSRE.2022.3210258

Lin, Y.-L., Potter-Baker, K. A., Cunningham, D. A., Li, M., Sankarasubramanian, V., Lee, J., Jones, S., Sakaie, K., Wang, X., Machado, A. G., & Plow, E. B. (2020). Stratifying chronic stroke patients based on the influence of contralesional motor cortices: An inter-hemispheric inhibition study. Clinical Neurophysiology : Official Journal of the International Federation of Clinical Neurophysiology, 131(10), 2516–2525. 10.1016/j.clinph.2020.06.016

Liu, K. P., Chan, C. C., Lee, T. M., & Hui-Chan, C. W. (2004). Mental imagery for promoting relearning for people after stroke: A randomized controlled trial. Archives of Physical Medicine and Rehabilitation, 85(9), 1403–1408. 10.1016/j.apmr.2003.12.035

Malouin, F., Richards, C. L., Jackson, P. L., Lafleur, M. F., Durand, A., & Doyon, J. (2007). The Kinesthetic and Visual Imagery Questionnaire (KVIQ) for assessing motor imagery in persons with physical disabilities: A reliability and construct validity study. Journal of Neurologic Physical Therapy: JNPT, 31(1), 20–29. 10.1097/01.NPT.0000260567.24122.64

Maris, E., & Oostenveld, R. (2007). Nonparametric statistical testing of EEG- and MEG-data. Journal of Neuroscience Methods, 164(1), 177–190. 10.1016/j.jneumeth.2007.03.024

mBrainTrain. (2025). *mBrainTrain software (Version 5.0.1.0)* [Computer software]. Belgrade, Serbia.

McFarland, D. J., Miner, L. A., Vaughan, T. M., & Wolpaw, J. R. (2000). Mu and beta rhythm topographies during motor imagery and actual movements. Brain Topography, 12(3), 177–186. 10.1023/A:1023437823106

Milde, C., Rance, M., Kirsch, P., Trojan, J., Fuchs, X., Foell, J., Bekrater-Bodmann, R., Flor, H., & Diers, M. (2015). Do mirror glasses have the same effect on brain activity as a mirror box? Evidence from a functional magnetic resonance imaging study with healthy subjects. PloS One, 10(5), e0127694. 10.1371/journal.pone.0127694

Morkisch, N., & Dohle, C. (2015). BeST - Berliner Spiegeltherapieprotokoll: Ein wissenschaftlich evaluiertes Manual zur Durchführung der Spiegeltherapie. Hippocampus Verlag. https://www.hippocampus.de/Best.60317.html?shl=1

Morkisch, N., Thieme, H., & Dohle, C. (2019). How to perform mirror therapy after stroke? Evidence from a meta-analysis. Restorative Neurology and Neuroscience, 37(5), 421–435. 10.3233/RNN-190935

Munzert, J., Blischke, K., & Krüger, B. (2015). Motor imagery of locomotion with an additional load: Actual load experience does not affect differences between physical and mental durations. Experimental Brain Research, 233(3), 809–816. 10.1007/s00221-014-4156-x

Munzert, J., Lorey, B., & Zentgraf, K. (2009). Cognitive motor processes: The role of motor imagery in the study of motor representations. Brain Research Reviews, 60(2), 306–326. 10.1016/j.brainresrev.2008.12.024

Neuper, C., & Pfurtscheller, G. (1999). Motor imagery and ERD. In G. Pfurtscheller & F. H. Lopes da Silva (Eds.), Event-Related Desynchronization: Handbook of Electroencephalography and Clinical Neurophysiology (Revised Edition Vol. 6, pp. 303–325). Elsevier.

Neuper, C., Scherer, R., Reiner, M., & Pfurtscheller, G. (2005). Imagery of motor actions: Differential effects of kinesthetic and visual-motor mode of imagery in single-trial EEG. Brain Research. Cognitive Brain Research, 25(3), 668–677. 10.1016/j.cogbrainres.2005.08.014

NICE. (2023). Stroke rehabilitation in adults. https://www.nice.org.uk/guidance/ng236/resources/stroke-rehabilitation-in-adults-pdf-66143899492549

Numata, K., Murayama, T., Takasugi, J., Monma, M., & Oga, M. (2013). Mirror observation of finger action enhances activity in anterior intraparietal sulcus: A functional magnetic resonance imaging study. Journal of the Japanese Physical Therapy Association = Rigaku Ryoho, 16(1), 1–6. 10.1298/jjpta.Vol16_001

Ono, T., Kimura, A., & Ushiba, J. (2013). Daily training with realistic visual feedback improves reproducibility of event-related desynchronisation following hand motor imagery. Clinical Neurophysiology: Official Journal of the International Federation of Clinical Neurophysiology, 124(9), 1779–1786. 10.1016/j.clinph.2013.03.006

Page, S. J., Levine, P., & Leonard, A. C. (2005). Effects of mental practice on affected limb use and function in chronic stroke. Archives of Physical Medicine and Rehabilitation, 86(3), 399–402. 10.1016/j.apmr.2004.10.002

Page, S. J., Levine, P., & Leonard, A. (2007). Mental practice in chronic stroke: Results of a randomized, placebo-controlled trial. Stroke, 38(4), 1293–1297. 10.1161/01.STR.0000260205.67348.2b

Page, S. J., & Peters, H. (2014). Mental practice: Applying motor PRACTICE and neuroplasticity principles to increase upper extremity function. Stroke, 45(11), 3454–3460. 10.1161/strokeaha.114.004313

Peirce, J., Gray, J. R., Simpson, S., MacAskill, M., Höchenberger, R., Sogo, H., Kastman, E., & Lindeløv, J. K. (2019). Psychopy2: Experiments in behavior made easy. Behavior Research Methods, 51(1), 195–203. 10.3758/s13428-018-01193-y

Pfurtscheller, G., Brunner, C., Schlögl, A., & Da Lopes Silva, F. H. (2006). Mu rhythm (de)synchronization and EEG single-trial classification of different motor imagery tasks. NeuroImage, 31(1), 153–159. 10.1016/j.neuroimage.2005.12.003

Pfurtscheller, G., & Lopes da Silva, F. H. (1999). Event-related EEG/MEG synchronization and desynchronization: Basic principles. Clinical Neurophysiology, 110(11), 1842–1857. 10.1016/S1388-2457(99)00141-8

Pfurtscheller, G., Neuper, C., & Krausz, G. (2000). Functional dissociation of lower and upper frequency mu rhythms in relation to voluntary limb movement. Clinical Neurophysiology, 111(10), 1873–1879. 10.1016/S1388-2457(00)00428-4

Picard, N., Matsuzaka, Y., & Strick, P. L. (2013). Extended practice of a motor skill is associated with reduced metabolic activity in M1. Nature Neuroscience, 16(9), 1340–1347. 10.1038/nn.3477

Pion-Tonachini, L., Kreutz-Delgado, K., & Makeig, S. (2019). Iclabel: An automated electroencephalographic independent component classifier, dataset, and website. NeuroImage, 198, 181–197. 10.1016/j.neuroimage.2019.05.026

Rothgangel, A., & Braun, S. (2013). Mirror Therapy: Practical Protocol for Stroke Rehabilitation. Pflaum Verlag. https://www.researchgate.net/publication/253235147_Mirror_Therapy_Practical_Protocol_for_Stroke_Rehabilitation 10.12855/ar.sb.mirrortherapy.e2013

Saleh, S., Adamovich, S. V., & Tunik, E. (2014). Mirrored feedback in chronic stroke: Recruitment and effective connectivity of ipsilesional sensorimotor networks. Neurorehabilitation and Neural Repair, 28(4), 344–354. 10.1177/1545968313513074

Saleh, S., Yarossi, M., Manuweera, T., Adamovich, S., & Tunik, E. (2017). Network interactions underlying mirror feedback in stroke: A dynamic causal modeling study. NeuroImage. Clinical, 13, 46–54. 10.1016/j.nicl.2016.11.012

Stagneth, L., Geritz, J., Hermann, G., Dahl, L., Zuber, L., Neumann, C., Lassen, J., Maetzler, C., Kranczioch, C., Welzel, J., & Maetzler, W. (2026). Eeg-based assessment of sustained attention in geriatric inpatients: A feasibility study. GeroScience. Advance online publication. 10.1007/s11357-026-02406-5

The MathWorks Inc. (2022). MATLAB version 9.12 (R2022a) [Computer software]. Natick, Massachusetts, United States. https://www.mathworks.com

The MathWorks Inc. (2022). MATLAB version 9.13 (R2022b) [Computer software]. Natick, Massachusetts, United States. https://www.mathworks.com

Thieme, H., Morkisch, N., Mehrholz, J., Pohl, M., Behrens, J., Borgetto, B., & Dohle, C. (2018). Mirror therapy for improving motor function after stroke. The Cochrane Database of Systematic Reviews, 7(7), CD008449. 10.1002/14651858.CD008449.pub3

Veale, J. F. (2014). Edinburgh Handedness Inventory - Short Form: A revised version based on confirmatory factor analysis. Laterality, 19(2), 164–177. 10.1080/1357650X.2013.783045

Villasana, F. C. (2022). Getting to know EEG artifacts and how to handle them in BrainVision Analyzer 2. Brain Products GmbH. https://pressrelease.brainproducts.com/eeg-artifacts-handling-in-analyzer/

Williams, S. E., Cumming, J., & Edwards, M. G. (2011). The functional equivalence between movement imagery, observation, and execution influences imagery ability. Research Quarterly for Exercise and Sport, 82(3), 555–564. 10.1080/02701367.2011.10599788

Wriessnegger, S. C., Brunner, C., & Müller-Putz, G. R. (2018). Frequency Specific Cortical Dynamics During Motor Imagery Are Influenced by Prior Physical Activity. Frontiers in Psychology, 9, 1976. 10.3389/fpsyg.2018.01976

Wriessnegger, S. C., Steyrl, D., Koschutnig, K., & Müller-Putz, G. R. (2014). Short time sports exercise boosts motor imagery patterns: Implications of mental practice in rehabilitation programs. Frontiers in Human Neuroscience, 8, 469. 10.3389/fnhum.2014.00469

Wright, D. J., Holmes, P., Di Russo, F., Loporto, M., & Smith, D. (2012). Reduced motor cortex activity during movement preparation following a period of motor skill practice. PloS One, 7(12), e51886. 10.1371/journal.pone.0051886

Yang, Y., Zhao, Q., Zhang, Y., Wu, Q., Jiang, X., & Cheng, G. (2018). Effect of Mirror Therapy on Recovery of Stroke Survivors: A Systematic Review and Network Meta-analysis. Neuroscience, 390, 318–336. 10.1016/j.neuroscience.2018.06.044

Yuan, H., Liu, T., Szarkowski, R., Rios, C., Ashe, J., & He, B. (2010). Negative covariation between task-related responses in alpha/beta-band activity and BOLD in human sensorimotor cortex: An EEG and fMRI study of motor imagery and movements. NeuroImage, 49(3), 2596–2606. 10.1016/j.neuroimage.2009.10.028

Zeng, W., Guo, Y., Wu, G., Liu, X., & Fang, Q [Qian] (2018). Mirror therapy for motor function of the upper extremity in patients with stroke: A meta-analysis. Journal of Rehabilitation Medicine, 50(1), 8–15. 10.2340/16501977-2287

Zhang, L., Qiu, F., Zhu, H., Xiang, M., & Zhou, L. (2019). Neural Efficiency and Acquired Motor Skills: An fMRI Study of Expert Athletes. Frontiers in Psychology, 10, 2752. 10.3389/fpsyg.2019.02752

Zhuang, J., Lei, X., Guo, X., Ding, L., & Jia, J. (2025). Motor and parietal cortex activity responses to mirror visual feedback in patients with subacute stroke: An EEG study. Clinical Neurophysiology Practice, 10, 12–21. 10.1016/j.cnp.2024.12.004

Zimmermann-Schlatter, A., Schuster, C., Puhan, M. A., Siekierka, E., & Steurer, J. (2008). Efficacy of motor imagery in post-stroke rehabilitation: A systematic review. Journal of Neuroengineering and Rehabilitation, 5, 8. 10.1186/1743-0003-5-8

