## Supplementary Material for "No Evidence of an Effect of a Single Mirror Visual Feedback Session on Subsequent Motor Imagery"

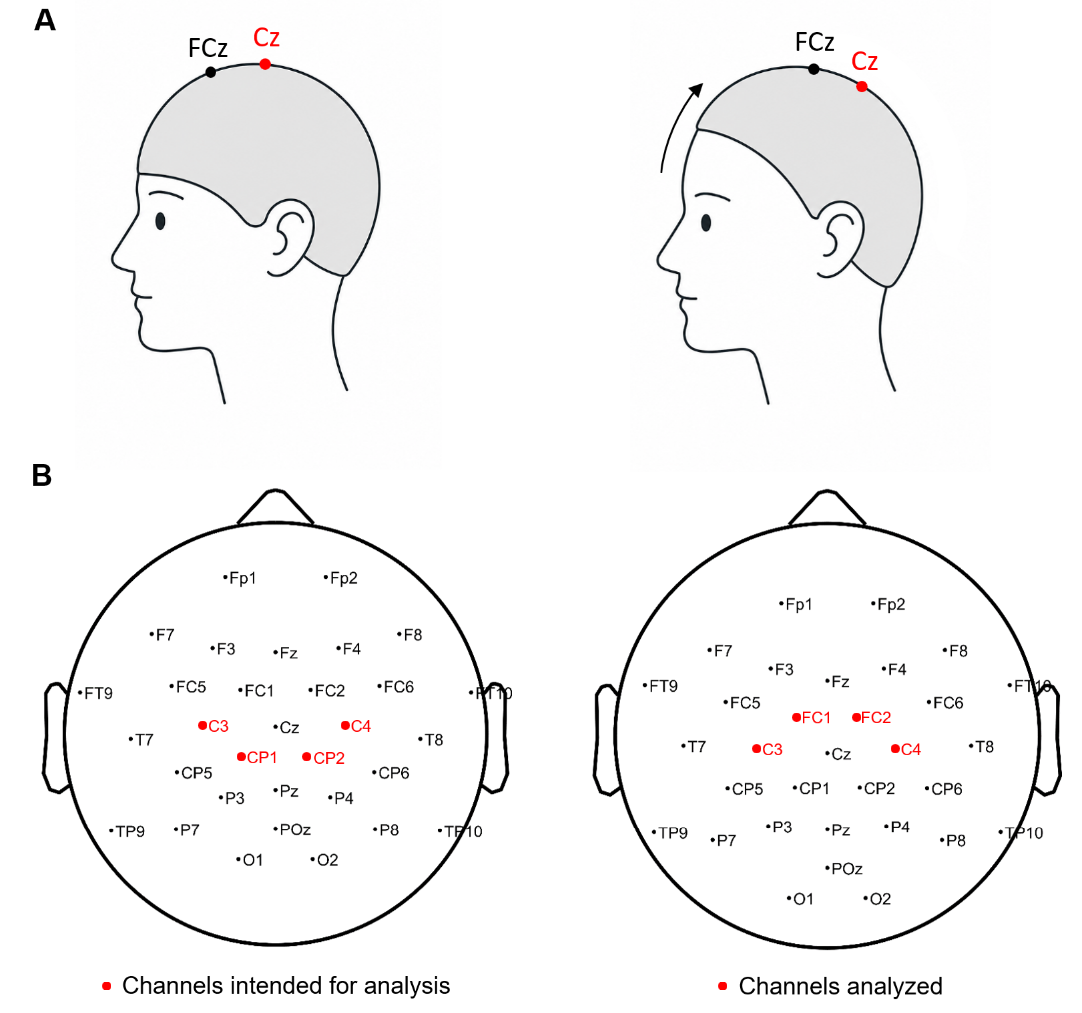

**Figure S1** | Systematic EEG cap displacement and channel selection. (A) Schematic illustration of the systematic posterior displacement of the EEG cap during data acquisition. The nominal reference electrode FCz was positioned at the 10-20 system anatomical Cz location, resulting in a posterior shift of the entire electrode montage. (B) Topographical illustration of the correctly aligned electrode configuration based on anatomical landmarks with the channels intended for analysis in red (left) and the shifted configuration with the channels selected for analysis in red (right). Channel selection in the shifted configuration was based on the digitized electrode positions and their spatial proximity to the intended sensorimotor coverage. Note that due to the cap displacement, nominal electrode labels in the shifted configuration (right) do not correspond to their locations in the 10-20 system. The illustrations in panel A were created using ChatGPT (OpenAI, accessed June 2026).

**Table S1** | Descriptive Statistics of MI Difficulty Ratings by MI Session Half

| MI Session | Half | Group | *M* | *SD* | Min | Max |
| --- | --- | --- | --- | --- | --- | --- |
| MI1 | First | MVF | 4.57 | 1.12 | 3.00 | 6.67 |
|  |  | Sham | 4.58 | 1.56 | 1.00 | 7.00 |
|  | Second | MVF | 4.72 | 0.89 | 3.33 | 6.00 |
|  |  | Sham | 4.53 | 1.45 | 1.67 | 6.67 |
| MI2 | First | MVF | 4.94 | 0.79 | 3.67 | 6.33 |
|  |  | Sham | 4.37 | 1.28 | 2.00 | 6.00 |
|  | Second | MVF | 4.76 | 0.86 | 3.33 | 7.00 |
|  |  | Sham | 4.17 | 1.24 | 2.33 | 6.00 |

Descriptive statistics of subjective motor imagery (MI) difficulty ratings averaged separately for the first and second half of MI1 and MI2. Ratings were obtained on a 7-point Likert scale (1 = very hard, 7 = very easy).

**Table S2** | Descriptive Statistics of Attention Ratings by Intervention Block

| Block | Group | *M* | *SD* | Min | Max |
| --- | --- | --- | --- | --- | --- |
| Block 1 | MVF | 6.44 | 0.86 | 4 | 7 |
|  | Sham | 6.25 | 0.85 | 4 | 7 |
| Block 2 | MVF | 6.33 | 1.14 | 3 | 7 |
|  | Sham | 6.05 | 0.95 | 4 | 7 |
| Block 3 | MVF | 6.44 | 0.92 | 4 | 7 |
|  | Sham | 6.20 | 0.95 | 4 | 7 |

Descriptive statistics of attention ratings collected during the intervention. Ratings were obtained on a 7-point Likert scale (1 = strongly disagree, 7 = strongly agree).

**Table S3** | Descriptive Statistics of Tiredness Ratings by Intervention Block

| Block | Group | *M* | *SD* | Min | Max |
| --- | --- | --- | --- | --- | --- |
| Block 1 | MVF | 1.94 | 1.11 | 1 | 4 |
|  | Sham | 2.35 | 1.31 | 1 | 5 |
| Block 2 | MVF | 1.89 | 1.08 | 1 | 4 |
|  | Sham | 2.05 | 1.05 | 1 | 4 |
| Block 3 | MVF | 1.94 | 1.06 | 1 | 4 |
|  | Sham | 2.40 | 1.05 | 1 | 4 |

Descriptive statistics of tiredness ratings collected during the intervention. Ratings were obtained on a 7-point Likert scale (1 = strongly disagree, 7 = strongly agree).

**Table S4** | Descriptive Statistics of Post-Procedure Questionnaire Items

| Item | Group | *M* | *SD* | Min | Max |
| --- | --- | --- | --- | --- | --- |
| Perceived difficulty of experiment | MVF | 3.17 | 1.20 | 1 | 5 |
|  | Sham | 3.30 | 1.63 | 1 | 6 |
| Clarity of instructions | MVF | 1.56 | 1.46 | 1 | 7 |
|  | Sham | 1.30 | 0.47 | 1 | 2 |
| Post-experimental physical state | MVF | 3.78 | 1.26 | 2 | 6 |
|  | Sham | 3.35 | 1.23 | 2 | 6 |

Descriptive statistics of post-procedure questionnaire items. Ratings were obtained on 7-point Likert scales. For Item 1, 1 = very easy, 7 = very hard; for Item 2, 1 = very clear, 7 = very unclear; for Item 3, 1 = very tired, 7 = very well rested.

**Table S5** | Mann-Whitney U Test Results for Post-Procedure Questionnaire Items

| Item | *U* | *p* | *r* |
| --- | --- | --- | --- |
| Perceived difficulty of experiment | 178.00 | .964 | 0.011 |
| Clarity of instructions | 172.00 | .775 | 0.044 |
| Post-experimental physical state | 215.00 | .296 | -0.194 |

r = rank biserial correlation.

**Table S6** | Results of the Bayesian Mixed-Design ANOVA on Mu ERD

| Effect | P(incl) | P(excl) | P(incl\|data) | P(excl\|data) | BF_incl_ |
| --- | --- | --- | --- | --- | --- |
| MI Session | 0.263 | 0.263 | 0.263 | 0.581 | 0.452 |
| Group | 0.263 | 0.263 | 0.281 | 0.495 | 0.568 |
| Hemisphere | 0.263 | 0.263 | 0.777 | 1.380×10^-4^ | 5629.018 |
| MI Session × Group | 0.263 | 0.263 | 0.092 | 0.142 | 0.647 |
| Hemisphere × Group | 0.263 | 0.263 | 0.153 | 0.351 | 0.435 |
| MI Session × Hemisphere | 0.263 | 0.263 | 0.080 | 0.337 | 0.237 |
| MI Session × Hemisphere × Group | 0.053 | 0.053 | 0.002 | 0.005 | 0.323 |

Results of a 2 × 2 × 2 bayesian mixed-design ANOVA with MI session and hemisphere as within-subject factors and group as the between-subject factor.

**Table S7** | Results of the Bayesian Mixed-Design ANOVA on Mu ERD Lateralization

| Effect | P(incl) | P(excl) | P(incl\|data) | P(excl\|data) | BF_incl_ |
| --- | --- | --- | --- | --- | --- |
| MI Session | 0.400 | 0.400 | 0.196 | 0.785 | 0.249 |
| Group | 0.400 | 0.400 | 0.261 | 0.720 | 0.362 |
| MI Session × Group | 0.200 | 0.200 | 0.019 | 0.049 | 0.386 |

Results of a 2 × 2 bayesian mixed-design ANOVA with MI session as the within-subject factor and group as the between-subject factor. Lateralization scores were calculated as the difference between left- and right-hemisphere mu ERD values (LH - RH), based on averages across the sensorimotor electrodes of interest within each hemisphere.

**Table S8** | Spearman Correlations Between Subjective Intervention Ratings and Changes in Contralateral Mu ERD

| Item | Group | Spearman's *ρ* | 95% CI | *p* |
| --- | --- | --- | --- | --- |
| Illusion realism | MVF | 0.44 | [−.06, .82] | .066 |
| Increasing realism | MVF | 0.04 | [−.44, .55] | .869 |
| Ownership (reversed) | MVF | −0.15 | [−.66, .38] | .558 |
| Left-arm movement | MVF | −0.34 | [−.80, .23] | .163 |
| Right-arm movement | MVF | −0.15 | [−.57, .34] | .561 |
| Bilateral movement | MVF | 0.06 | [−.47, .59] | .826 |
| Task demand | MVF | 0.20 | [−.31, .60] | .431 |
| Attention | MVF | 0.27 | [−.28, .70] | .281 |
| Tiredness | MVF | 0.11 | [−.43, .60] | .677 |
| Task demand | Sham | −0.19 | [−.65, .31] | .434 |
| Attention | Sham | 0.21 | [−.29, .63] | .373 |
| Tiredness | Sham | −0.42 | [−.70, .01] | .063 |

Spearman correlation coefficients (*ρ*) and corresponding 95% bootstrap confidence intervals are reported. Confidence intervals were estimated using 5,000 bootstrap samples. Ratings were obtained on 7-point Likert scales. For Items 1-3 and 7-12, 1 = strongly disagree, 7 = strongly agree; for Items 4-6, 1 = never, 7 = always. MVF = mirror visual feedback.

**Table S9** | Spearman Correlations Between Subjective Intervention Ratings and Changes in Mu ERD Lateralization

| Item | Group | Spearman's *ρ* | 95% CI | *p* |
| --- | --- | --- | --- | --- |
| Illusion realism | MVF | -.02 | [-.46, .45] | .950 |
| Increasing realism | MVF | -.13 | [-.59, .42] | .594 |
| Ownership (reversed) | MVF | .23 | [-.24, .67] | .359 |
| Left-arm movement | MVF | .03 | [-.43, .50] | .919 |
| Right-arm movement | MVF | .09 | [-.44, .63] | .728 |
| Bilateral movement | MVF | -.31 | [-.73, .20] | .215 |
| Task demand | MVF | .11 | [-.42, .59] | .675 |
| Attention | MVF | -.45 | [-.82, .07] | .059 |
| Tiredness | MVF | .08 | [-.42, .55] | .759 |
| Task demand | Sham | -.30 | [-.65, .14] | .205 |
| Attention | Sham | .43 | [.06, .67] | .061 |
| Tiredness | Sham | -.38 | [-.75, .08] | .102 |

Spearman correlation coefficients (*ρ*) and corresponding 95% bootstrap confidence intervals are reported. Confidence intervals were estimated using 5,000 bootstrap samples. Ratings were obtained on 7-point Likert scales. For Items 1-3 and 7-12, 1 = strongly disagree, 7 = strongly agree; for Items 4-6, 1 = never, 7 = always. MVF = mirror visual feedback.
